# Representational Dynamics Preceding Conscious Access

**DOI:** 10.1101/2020.08.30.274019

**Authors:** Josipa Alilović, Dirk van Moorselaar, Marcel Graetz, Simon van Gaal, Heleen A. Slagter

## Abstract

Our senses are continuously bombarded with more information than our brain can process up to the level of awareness. The present study aimed to enhance understanding on how attentional selection shapes conscious access under conditions of rapidly changing input. Using an attention task, EEG, and multivariate decoding of individual target- and distractor-defining features, we specifically examined dynamic changes in the representation of targets and distractors as a function of conscious access and the task-relevance (target or distractor) of the preceding item in the RSVP stream. At the behavioral level, replicating previous work and suggestive of a flexible gating mechanism, we found a significant impairment in conscious access to targets (T2) that were preceded by a target (T1) followed by one or two distractors (i.e., the attentional blink), but striking facilitation of conscious access to targets shown directly after another target (i.e., lag-1 sparing and blink reversal). At the neural level, conscious access to T2 was associated with enhanced early- and late-stage T1 representations and enhanced late-stage D1 representations, and interestingly, could be predicted based on the pattern of EEG activation well before T1 was presented. Yet, across task conditions, we did not find convincing evidence for the notion that conscious access is affected by rapid top-down selection-related modulations of the strength of early sensory representations induced by the preceding visual event. These results cannot easily be explained by existing accounts of how attentional selection shapes conscious access under rapidly changing input conditions, and have important implications for theories of the attentional blink and consciousness more generally.

## 1 Introduction

Over the past few decades, research has shown that visual information processing preceding conscious access tends to cluster in several functionally distinct stages after stimulus presentation (Carlson, Tovar, Alink, & Kriegeskorte, 2013; Grootswagers, Robinson, & Carlson, 2019; Kaiser, Oosterhof, & Peelen, 2016; Marti & Dehaene, 2017; Marti, Sigman, & Dehaene, 2012; Sergent, Baillet, & Dehaene, 2005; Weaver, Fahrenfort, Belopolsky, & Van Gaal, 2019). The early and intermediate phases of stimulus processing, up to ~300 ms after stimulus presentation, are characterized by bottom-up and local recurrent processing in sensory cortex (Dehaene, Changeux, Naccache, Sackur, & Sergent, 2006; Lamme & Roelfsema, 2000). During these stages, stimulus processing is primarily bottom-up and non-conscious, supported by a greatly parallel processing architecture, which permits multiple visual stimuli to be represented in the brain at the same time. The subsequent processing phase is however selective to those stimuli amplified in a top-down manner depending on their goal relevance, i.e., that are attentionally selected (Marti & Dehaene, 2017; Olivers & Meeter, 2008; Sergent et al., 2005; Sigman & Dehaene, 2008). At this relatively late processing stage, stimuli are encoded into working memory, a process thought critical for translating fleeting sensory representations into a more durable consciously accessible format (Dehaene et al., 2006; Marti & Dehaene, 2017; Olivers & Meeter, 2008). Yet, the current literature accommodates two conflicting views about what determines whether or not a stimulus becomes available for conscious access: the serial nature of limited-capacity late-stage processing, or dynamic gating effects on lower-stage sensory representations.

Limited-capacity models (Chun & Potter, 1995; Jolicœur & Dell’Acqua, 1998; for review see Dux & Marois, 2009 and Olivers & Meeter, 2008) and some theories of consciousness (e.g. Global Neuronal Workspace; Dehaene, Charles, King, & Marti, 2014) postulate a serial bottleneck during late-stage processing. Specifically, only one item can be encoded at a time by late-stage capacity-limited processes allowing conscious access (Marti & Dehaene, 2017; Marti et al., 2012; Sergent et al., 2005; Sigman & Dehaene, 2008). According to these accounts, conscious access fails when an item cannot be attentionally selected for late-stage encoding, for example when this stage is still occupied by a previous item. This is well illustrated by the so-called attentional blink (AB): an impairment in identifying a second target (T2) presented after a first target (T1) within close temporal proximity (200 to 500 ms) in a rapid stream of distractor stimuli (Raymond, Shapiro, & Arnell, 1992). According to limited-capacity accounts, conscious access to T2 fails because T1 encoding into working memory ties up limited processing resources, rendering them temporarily unavailable for T2 (Lagroix, Spalek, Wyble, Jannati, & Di Lollo, 2012; Marti & Dehaene, 2017; Marti, Sigman, & Dehaene, 2012; Sergent, Baillet, & Dehaene, 2005).

Notwithstanding their popularity, limited-capacity accounts fall short in explaining several more recent behavioral observations. First, overall high target accuracy is observed, even for targets presented in the typical AB time window, when targets are presented sequentially with no intervening distractors (e.g., TTTDD; T – target; D – distractor), a phenomenon called sparing (Di Lollo, Kawahara, Ghorashi, & Enns, 2005; Lunau & Olivers, 2010; Olivers, Hilkenmeier, & Scharlau, 2011; Olivers, Van Der Stigchel, & Hulleman, 2007). What is more, T2 performance often exceeds T1 performance when the two targets are shown consecutively (Dell’Acqua, Doro, Dux, & Losier, 2016; Di Lollo et al., 2005; Olivers et al., 2011). Even more problematic for limited-capacity accounts is the so-called AB reversal, whereby in a TDTT sequence T3 seems to “escape” the AB. That is, T3 accuracy is higher when T3 is preceded by a target (TDTT) than a distractor (TTDT) and higher than T2 accuracy at this same temporal position in the stream (TDDT) (Kawahara, Kumada, & Di Lollo, 2006; Olivers et al., 2007). These findings are difficult to explain assuming a T1-triggered late-stage bottleneck.

In an alternative account, the boost and bounce theory of temporal attention (Olivers & Meeter, 2008), the AB, sparing of conscious access, and blink reversal are consequences of (dys)functional gating of information into working memory. More specifically, this theory proposes that a combination of excitatory and inhibitory gate neurons form an attentional gating system into working memory, i.e., implement the attentional set, and provide excitatory (“boost”) and inhibitory (“bounce”) feedback upon target and distractor detection, respectively. Critically, this top-down feedback peaks rapidly, approximately 100 ms after stimulus presentation (e.g. Shimozaki, Chen, Abbey, & Eckstein, 2007; Wyble, Bowman, & Potter, 2009, for a review, see Olivers, 2012) thereby also affecting the chance of conscious access for the following item. In this account, the attentional blink to T2 is caused by strong inhibitory feedback (a bounce) triggered by the distractor after T1 (D1), that itself was accidentally boosted by strong excitatory feedback evoked by T1. This account can also readily explain sparing: if the first post-T1 stimulus in the stimulus stream is T2, this stimulus, as well as other immediately ensuing target stimuli, will be boosted into working memory (hence the observation of extended sparing in a TTTDD sequence). Rapid reversal of the AB is similarly explained by the workings of this rapid gating system: T3 is relatively boosted when it directly follows T2 (TDTT) compared to a distractor (TTDT), rendering it more likely that it will gain access to consciousness. Thus, according to this account, the attentional blink reflects dysfunctional gating of information to late-stage processing and not a capacity limitation of late stage visual information processing per se. Another influential model, the serial token/simultaneous type (STST/eSTST) model, also attributes the AB to dysfunctional gating of information into working memory in that during T1 encoding, an attentional ‘blaster’ is temporally unavailable to boost the representation of next task-relevant items (Bowman & Wyble, 2007; Wyble, Bowman, & Nieuwenstein, 2009). In this model, a T1-triggered attentional enhancement can also strengthen the representation of a subsequently presented item, such as D1. Yet, this model does not assign a critical role to D1, as the AB is caused by T1-triggered resource depletion (i.e., the unavailability of the blaster during T1 encoding), not enhanced D1 processing.

Neural evidence for rapid boost and/or bounce gating of conscious access is so far scarce and relatively inconsistent. ERP studies have reported an attentional selection response to T1, the frontal selection positivity component peaking approximately 250 ms after the onset of T1, followed 100-150ms later by a frontal negativity, on target-present (TD) in contrast to target-absent (DD) trials (Martens, Munneke, Smid, & Johnson, 2006). The frontal negativity was furthermore found to increase in amplitude as the number of stimuli that had to be ignored grew, which was also related to a deficit in awareness of the subsequent target, hence presumably signaling stronger frontal gating or inhibition (Niedeggen, Hesselmann, Sahraie, Milders, & Blakemore, 2004). These findings were interpreted as post-T1 attentional enhancement followed by distractor-triggered inhibition, and taken as evidence for the boost and bounce theory (Olivers & Meeter, 2008). However, a more recent study that compared the negativity arising after the frontal selection positivity component between two conditions that differed only in the temporal position of the first post-T1 distractor (TD and TTD) did not observe a latency shift of the frontal negativity component, challenging the assumption that the frontal negativity reflects a distractor-evoked inhibitory response (Dell’Acqua et al., 2016). Furthermore, a recent study that used inverted encoding modeling to decode the orientation of each item in the AB task (stimuli were oriented gratings with different spatial frequencies), which could thus isolate single-item processing dynamics, only observed AB-related changes in early orientation tuning to T2, but no differences in the representational strength of D1 as a function of T2 visibility (Tang et al., 2020). The lack of effects on the sensory representation of D1 in this study may argue against the notion that accidental selection or boosting of D1 causes the AB to T2. However, no differences in T1 representation were observed either, which is surprising as limited capacity accounts, the STST/eSTST model, and boost and bounce theory all predict that the attentional blink is related to having to encode T1, albeit only indirectly in the latter account. Possibly, as only spatial frequency but not orientation was a predictable/defining feature of targets in the Tang et al. study, and hence only spatial frequency could drive attentional search, their orientation decoding may have been less sensitive to top-down feature selection-related effects.

The present EEG study aimed to advance understanding of how attentional selection shapes conscious access using multivariate decoding of individual target- and distractor-defining features, and an attention task that allowed us to examine dynamic changes in the representation of individual targets and distractors in attentional blink, sparing and AB reversal conditions. Specifically, we tested two predictions that disentangle limited-capacity and boost and bounce accounts at the neural level. First, while limited-capacity accounts generally assign no critical role to D1, the boost and bounce theory posits that the AB is related to accidental selection of D1 for late-stage processing: T1-evoked top-down feedback meant to strengthen its low-level neural representation also accidentally amplifies the sensory representation of D1, because of its close temporal proximity to T1, boosting it into working memory. Therefore, this account predicts quantifiable differences in the quality of D1 representations in T2 seen vs. unseen trials, which we examined here directly. Second, as noted above, limited-capacity accounts have trouble explaining the behavioral observations of extended sparing and AB reversal, which the boost and bounce account links to dynamic attentional gating of information, depending on the nature of the preceding item in the stream. Here, we hence also investigated changes in the quality of target (e.g., T3) and distractor representations as a function of the nature of the preceding item in the stream (i.e., target or distractor).

To test these predictions, participants performed an attention task (cf. Olivers et al., 2007) in which they had to identify up to three target numbers presented in a rapid serial visual presentation (RSVP) stream of distractor letters, while in each trial, we varied the number of targets and their temporal order. Concurrently, we measured their brain activity using EEG to which we applied multivariate pattern analysis (MVPA) (Grootswagers, Wardle, & Carlson, 2017; King & Dehaene, 2014). This approach enabled us to identify individual stimulus-specific sensory representations at distinct processing stages with high temporal precision and at the whole-brain level. MVPA neural pattern classifiers trained at each time point were also applied to all other time points, so that using the resulting generalization across time matrix, we could also examine whether and when neural patterns were stable and thus generalized across time (King & Dehaene, 2014). Recent studies have identified (at least) three visual information processing stages that can be separated using the generalization across time approach. Its early diagonal portion (<200 ms) is thought to reflect early-stage sensory processes driven by bottom-up input characteristics (Fahrenfort et al., 2017; King et al., 2016; Marti & Dehaene, 2017), and a late-stage (~300-600 ms) period with a sustained temporal profile, extending off diagonal, that correlates with conscious access and task-related goals (Marti & Dehaene, 2017; Meijs et al., 2019; Weaver et al., 2019). Early decoding can also extend off the diagonal, reflecting maintenance of a low-level sensory representation over time (Meijs et al., 2019; Weaver et al., 2019). Building on this body of work, we specifically examined dynamical changes in neural representations across these different processing stages, with the ultimate goal of gaining a better understanding of the underlying processing architecture that determines conscious access.

## 2 Methods

### 2.1 Participants

Thirty-five right-handed subjects (29 female, mean age = 20.91 years, SD=2.16 years), all students from the University of Amsterdam, who reported normal or corrected-to-normal vision and no history of a psychiatric or neurological disorder, participated in this study. Participants gave written informed consent prior to the start of the study and received research credits or money (10 euros per hour) for their participation. The study was approved by the ethical committee of the Department of Psychology of the University of Amsterdam. One participant was excluded from the final analyses because of misunderstanding the task instructions, while two other participants dropped out before finishing the third session. The final sample thus consisted of thirty-two participants who each completed three EEG sessions (27 female, mean age = 20.78 years, SD=1.83 years).

### 2.2 Stimuli and apparatus

All stimuli were generated using Matlab 8 and Psychtoolbox-3 software (Kleiner et al. 2007) within a Matlab environment (Mathworks, RRID:SCR_001622). Stimuli were presented on a 1920×1080 pixels BenQ XL2420Z LED monitor at a 120-Hz refresh rate on a “black” (RGB: [0 0 0], ± 3 cd/m^2^) background and were viewed with a distance of 90 cm from the monitor.

### 2.3 Procedure

The study consisted of three EEG sessions in which participants either searched for one target in an RSVP stream (localizer task session) or for up to three targets in an RSVP stream (two attention task sessions), while their brain activity was recorded using EEG. EEG cap placement was standardized for each participant across the three recording sessions to reduce the chance that differences in electrode locations across sessions contributed to our decoding results. Specifically, in each session, we measured the distance from the nasion to the inion, across the top of the head, assuring that the central Cz electrode was positioned exactly in the middle. We then measured the distance from the tragus of the left ear to the tragus of the right ear, across the top of the head, again making sure that the Cz was located in the middle.

### 2.4 Localizer task

The study started with one 180-minute localizer task session in which, on each trial, participants had to identify a single target embedded in an RSVP stream of 13 distractor stimuli. In half of the trials, the target was a number presented among distractor letters, while in the other half of trials, the single target was a letter presented among distractor numbers. The target stimulus, one of eight numbers (2-9) or one of eight letters (A, D, H, K, L, M, R, U), always appeared on positions 5-9 (balanced across trials). The distractor stream consisted of the eight stimuli of the other category, presented in a random fashion without consecutive repetitions. All stimuli were shown at fixation in a monospaced font (font size: 55 points) in white (RGB: [255 255 255]) for 83 ms with no inter-stimulus interval (ISI). Each trial started with a fixation cross for 400ms +/− 150ms jitter (25ms step size). After the last stimulus in the stream, the fixation cross was shown again for another 600 ms, after which participants were asked to identify which target number or letter (depending on the block) they had seen using 8 yellow-marked keys (a, s, d, f, j, k, l and;) on the keyboard in front of them, which spatially corresponded to 8 numbers or letters shown on the computer screen in a specific order, for example: 5 6 7 8 9 2 3 4. In this example, if for instance they saw the target number 7, they needed to press the third yellow key on the keyboard. The position of items on the screen and hence their associated response key varied across trials (e.g., 2 3 4 5 6 7 8 9; 4 5 6 7 8 9 2 3; etc.). This was done so that our subsequent decoding analysis could not pick up on any consistent stimulus-response relationships and decoding results would not be confounded by activity related to specific response preparation. Numbers and letters were always presented in ascending order, with the starting item varying from trial to trial. For instance, on ⅛ of trials the response sequence started with the number 2, on other ⅛ of trials with the number 3 and so on. This resulted in eight possible number response orders and eight possible letter response orders, which were presented equally often over the course of the experiment. Participants were asked to maintain fixation at all times except, if necessary, during the response period.

The task consisted of 1440 trials, presented in 18 blocks. Half of the participants first completed 9 blocks of trials in which they needed to identify a target letter, while in the remaining 9 blocks they needed to identify a target number. The order of letter and number blocks was reversed for the other half of the participants. Blocks of trials were interleaved with self-paced breaks, except after every forth block when a longer break, paced by the experimenter, was administered. After every block, participants received feedback about their performance (percentage of correct target identifications for that block).

EEG activity was concurrently recorded so that we could build classifiers to decode identity-specific neural representations of the different target stimuli, unbiased by any task manipulation that we employed in the following two experimental sessions (see below). The EEG data was used to build two types of classifiers: one that classified eight different letters and one that classified eight different numbers.

### 2.5 Attention task

In the second and the third session of the study, participants performed an attention task (adopted from Olivers et al., 2007), while their brain activity was again recorded using EEG. On each trial, they saw 1-3 target numbers (T1, T2, T3) embedded in a stream of distractor letters. Participants’ task was to report the identity of all targets they had seen in the stimulus stream.

Stimuli and the design of the attention task were identical to the localizer task except for the following differences. Each RSVP stream consisted of 18 stimuli in total. Target stimuli were numbers ranging from 2 to 9, while distractors could be 15-17 letters (A, D, H, K, L, M, R, U, C, E, F, G, I, J, N, O, P, T, V, W, X, Y, Q, Z). Each number appeared as T1, T2 and T3 equally often. Only the distractor letters A, D, H, K, L, M, R, U, shown as targets in the localizer task, could appear at positions 5 to 9. We pseudo-randomized their order such that each letter appeared at each given position within a condition equally often. The other distractor letters were randomly presented at the other temporal positions (i.e., 1-4, 10-17). The same target number and distractor letter was never repeated within a trial.

Each trial started with a fixation cross shown at the center of the screen for 700 ± 100 ms with a 25 ms step size. After the stream ended, the fixation cross reappeared for 800 ms, after which the response screen appeared. The manner of responding was identical to the localizer task, except that participants could now report more than one target. They were instructed to report any target seen and in case of multiple targets, in the order they had seen them in the stream, but the latter was not emphasized as crucial. After indicating seen targets, participants needed to press the spacebar to confirm their entry and to start the next trial. Participants were asked to maintain fixation at all times except, if necessary, during the response period.

We manipulated the number of targets shown in each trial (1-T: 12%, 2-T: 50%, 3-T: 38% of trials) and the lag at which T2 and T3 were presented after T1 similar to Olivers et al. (2007) to be able to measure four critical phenomena: the attentional blink, lag-1 sparing, extended sparing, and AB reversal. Specifically, a combination of the two factors (number of targets and lag) resulted in following 8 conditions: 1. T_1_D_1_D_2_D_3_..T_2_ (12% long-lag trials), 2. T_1_T_2_D_1_D_2_D_3_ (10% lag-1 trials), 3. T_1_D_1_T_2_D_2_D_3_ (14% lag-2 trials), 4. T_1_D_1_D_2_T_2_D_3_ (14% lag-3 trials), 5. T_1_T_2_T_3_D_1_D_2_ (10% extended sparing trials), 6. T_1_T_2_D_1_T_3_D_2_ (14% lag-2 and 4 trials), 7.T_1_D_1_T_2_T_3_D_2_ (14% AB reversal trials), 8. D_1_D_2_D_3_D_4_T_1_ (12%, single target trials). T1 was shown in each trial at temporal position 5 in the stream in all conditions, except for the condition with one target (1-T condition: 8), when a single target was presented at one of the late temporal positions (13-16). In conditions with two targets (2-T conditions: 2, 3, 4), T1 was followed by T2 at lags 1, 2 or 3, or at one of the long lags 8, 9, 10 or 11 (in long-lag trials). In the three conditions with three targets (T-3 conditions: 5, 6, 7), T1 was followed by T2 either at lag 1 or 2, whereas T3 appeared either at lag 2 or 3 (depending on T2).

In each of the two 180-minute sessions, each participant completed 16 blocks of 67 trials. Between blocks, participants could take a short break. After every fourth block, there was an enforced, longer break. After each block, participants received feedback about their performance (percentage of correct T1 identification). The experimenter also kept track of the percentage of T2 and T3 false alarms and warned participants not to guess if their false alarm rate exceeded 20%.

### 2.6 EEG recording and preprocessing

During each session, participants’ brain signals were sampled continuously at 512 Hz using a BioSemi ActiveTwo system (www.biosemi.com) with 64 scalp electrodes placed according to the 10/10 system. Two electrodes were placed on the earlobes for offline rereferencing and four electrooculographic (EOG) electrodes measured horizontal and vertical eye movements. After data acquisition, preprocessing and subsequent analyses were performed using custom-written analysis scripts which are publicly available and can be downloaded at https://github.com/dvanmoorselaar/DvM. These custom written analysis scripts are largely based on MNE software functionalities (Gramfort et al., 2014). EEG data were referenced offline to the average activity recorded at the earlobes and high-pass filtered using a zero-phase ‘firwin’ filter at 0.1 Hz as implemented in MNE to remove slow drifts. EEG signals were visually inspected for extremely noisy or malfunctioning electrodes, which were temporarily removed from subsequent preprocessing (20 participants had no channels removed, while the median=2 (range=2) for the remaining 12 participants). Epochs with excessive EMG artifacts were rejected using an adapted version of the *ft_artifact_zvalue* automatic trial rejection procedure, as implemented in the Fieldtrip toolbox (Oostenveld, Fries, Maris, & Schoffelen, 2011, http://fieldtriptoolbox.org). This function applies a frequency filter between 110 and 140 Hz and assigns a variable z-value score cutoff per participant based on the within-subject variance of z scores (cf. van Moorselaar & Slagter, 2019). On average, 16.3%, 15.9% and 17% of trials were removed per participant in the first, second and third session, respectively, using this approach.

Epochs of EEG data containing all events of interest for a given trial were created for the localizer and the attention task data from −400 to 1440 ms and −400 to 2000 ms, respectively, centered on T1 presentation time. Epoched data was baseline corrected to the average activity between −200 and 0ms pre-T1 stimulus presentation. Independent component analysis (ICA), as implemented in MNE using the ‘extended-infomax’ method, was performed on non-epoched 1 Hz high pass-filtered data to remove eye-blink components from the 0.1 Hz filtered data (cf. van Moorselaar & Slagter, 2019). Components topographies were visually inspected and compared to EOG signals. A single eye blink component per session was removed from epoched participant’s EEG data. Malfunctioning electrodes were then interpolated using spherical splines (Perrin, Perring, Bertrand, & Echallier, 1989).

### 2.7 Multivariate decoding analyses

Multivariate pattern analysis (MVPA) was applied to EEG data to decode patterns of neural activity specific to each target number and each distractor letter (i.e., only those shown on positions 5-9) in the RSVP streams for each condition of interest in the attention task. Classifiers were trained on the localizer task data and applied to the attention task data (cross-task decoding). This allowed us to examine if 1) the strength of target and distractor stimulus-specific representations preceding T2 were associated with conscious access to T2, and 2) whether stimulus-specific representations were generally stronger versus weaker depending on whether they were shown on boosted or bounced positions in the RSVP stream.

In order to decrease the computational time needed for MVPA, we downsampled the EEG data to 128Hz and shortened epochs used for training and testing classifiers to −200 to 900 ms with respect to the presentation time of the stimulus of interest. Decoding analyses were applied using the Scikit-learn Python (Python Software Foundation, https://www.python.org/) package. We applied a linear discriminant analysis using default settings (Pedregosa, Weiss, & Brucher, 2011) to raw EEG data recorded at all 64 electrodes, using each time sample in the cross-task validation procedure or 10-fold cross-validation procedure (see below). When classifiers were trained and tested on each time sample of two independent datasets to decode classes of stimuli, training was done using the localizer task and testing was done on the attention task data. Based on the localizer task data, we thus built letter-specific and number-specific classifiers for each time point of the data, which were then applied to the attention task data. The multi-class decoding problem (i.e. decoding 8 different numbers and 8 different letters) was formulated as multiple binary classification problems such that each class was tested against all other classes (i.e. the so-called “one-vs-all” approach) (Bishop, 2006). This means that a single classifier is trained per class to decode that class from the “other class”, consisting of *all* remaining classes (e.g., the number two versus any other possible number). This is done serially for each class, i.e., for each of eight letters or eight numbers shown in the localizer task. Each classifier is then applied to an unseen sample from the testing set of the attention task, for which the label is predicted by choosing the classifier that yields the highest confidence score for that class. The final score is obtained by averaging scores for all classes.

We also used the localizer task or attention task only in combination with a 10-fold cross-validation procedure in order to within-task decode target and distractor stimulus-specific representations (multi-class decoding) and target versus distractor stimulus classes (binary decoding, “target” vs. “distractor”), respectively. One participant’s data was not included in the analysis of the attention task when decoding T2 stimulus-specific representation due to an insufficient number of trials in a fold to train the classifier on all possible T2 numbers. Using the 10-fold cross-validation scheme we also decoded whether T2s were reported seen versus whether they were missed in the attention task, using “seen” vs. “unseen” labels for decoding. In general, in the 10-fold cross-validation scheme, the classifier was trained on 90% of the data to classify between stimulus classes, and then tested on the remaining 10% of the data. This procedure was repeated 10 times, until all data were tested exactly once. The percentage of correct class assignments was averaged across the 10 folds. Classifier’s performance in separating two or more classes of stimuli was expressed as the area under the curve (AUC), which indicates the degree of separability between classes by integrating the receiver operating characteristic (ROC) curve (Fawcett, 2006; Myerson, Green, & Warusawitharana, 2001). The training procedure was done on balanced stimulus classes, which means that each stimulus class was present equally often during training.

We used the so-called generalization across time approach in applying the pattern classifiers (King & Dehaene, 2014) - a classifier trained on a specific time point was tested on that time point as well as on all other time points. The resulting generalization across time matrix (training time on y-axis x testing time on x-axis) for targets and distractors can therefore reveal periods during which a representation is stable, i.e. generalizes across time. For instance, a classifier trained to distinguish between stimulus classes at 170 ms can be applied to an entire time course or smaller segments of time data (e.g., 170-220 ms and 300-600 ms) to test whether a stimulus representation is maintained. This approach is thus informative of stimulus-specific representations at different stages of visual information processing, permitting us to examine when in time and at what processing stage representations might be modulated, and comparing the results to predictions from the two theoretical account.

### 2.8 ERP analyses

Awareness of stimuli such that they can be reported is typically associated with a late (300-500ms) broadly distributed positive P3 ERP component (Cohen, Ortego, Kyroudis, & Pitts, 2020; Dehaene & Changeux, 2011; Derda et al., 2019; Sigman & Dehaene, 2008). For example, it has been shown that only seen T2s elicit a P3 (Vogel, Luck, & Shapiro, 1998). Here, we also aimed to replicate this finding. To this end, we selected a subset of centro-parietal channels (POz, Pz, CPz, CP1, CP2, P1, P2, PO3, PO4) which are known to capture the P3 component topography and created ERP waveforms using trials in which T1 was correctly identified, but splitting the analysis on correctly identified T2s (i.e. allowing order reversals in report, which meant that a response was considered correct when a correct number was reported at the end of a trial irrespective of the report order) and missed or incorrectly identified T2s (T2 seen and unseen in further text) in the T_1_D_1_T_2_D_2_D_3_ condition. We also computed the P3 to correctly-identified T1s using the T_1_D_1_D_2_D_3_..T_2_ condition in which T2s were shown at late latencies and could thus not impact the T1-elicited P3 component. By contrasting ERP waveforms to T2-late, T2-seen and T2-unseen trials, we could thus better distinguish between P3’s elicited by T1 and T2 stimuli. All ERP waveforms were time-locked to T1 presentation time.

### 2.9 Statistical analyses

#### 2.9.1 Behavior

To evaluate behavioral performance, for each participant we computed the percentage of correct target identifications in the localizer and attention task. In the attention task, given that participants could report up to three targets, percentage correct for each target, i.e. separately for T1, T2 and T3, was computed by taking into account the total number of trials in which that target was present. As in Olivers et al. (2007), we computed percentages of correct target identifications in the attention task for each target separately allowing order reversals in report. This means that a response was considered correct when the correct number was reported at the end of the trial even if the report order did not match the target presentation order. Furthermore, as in Olivers et al., accuracy for the post-T1 targets was contingent on T1 correct identification. For the attention task, we also removed trials which were rejected from the EEG dataset during preprocessing using automatic trial rejection procedure.

To verify the presence of an attentional blink, sparing, and blink reversal, we conducted three separate repeated-measures ANOVAs as in Olivers et al. (2007), with T1, T2 and/or T3 identification accuracies as the dependent variable. Note that we included temporal position (TP) instead of lag as a within-subject factor in these statistical analyses to denote the timing of an event in the stream. This is because our ANOVA models could include T1 performance as well. At the earliest, T1 could appear on position 5 in the stream, which we coded as TP1 into the ANOVA analysis. Accordingly, targets on position 9 in the stream, for instance, were coded as TP5 targets. Moreover, in order to evaluate the performance for T1s and T2s shown on the 4 late positions in the single-target and long lag conditions (conditions 1 and 8), respectively, we aggregated performance accuracies across those positions within a condition and entered the score as the “late TP” target. One omnibus repeated-measures ANOVA was not possible because not all conditions had targets at same temporal positions. To verify the presence of the AB, we first conducted a one-way repeated measures ANOVA with T2 identification accuracy obtained in 2-T conditions (i.e., T_1_T_2_D_1_D_2_D_3_, T_1_D_1_T_2_D_2_D_3_, T_1_D_1_D_2_T_2_D_3_, and T_1_..D_1_D_2_D_3_T_2_) as the dependent variable and Temporal Position (TP 2, 3, 4, and late (13-16)) as a within subjects factor. To determine evidence for extended sparing (Di Lollo et al., 2005; Olivers et al., 2007), we conducted a repeated measures ANOVA with Number of Targets (2-T or 3-T) and Temporal Position (TP 1-3) as within subject factors based on target accuracy in the 2-T conditions (T_1_T_2_D_1_D_2_D_3_ and T_1_D_1_T_2_D_2_D_3_) and the 3-T condition (T_1_T_2_T_3_D_1_D_2_). Finally, we statistically verified the presence of attentional blink reversal using a repeated measures ANOVA with Number of Targets (2-T vs. 3-T) and Temporal Position (TP 1, 3 and 4) as within subject factors based on target accuracy in the following conditions: T_1_D_1_T_2_D_2_D_3_, T_1_D_1_D_2_T_2_D_3_, and T_1_D_1_T_2_T_3_D_2_. In all analyses, significant main and interaction effects were followed-up by paired-sample t-tests.

#### 2.9.2 EEG

In order to statistically evaluate classifier’s performance across time in picking up stimulus-specific representations, we tested whether classifier’s performance (AUC) at each time point of the generalization across time matrix was significantly different than at chance decoding. For this, we applied group-level permutation testing with cluster correction for multiple comparisons (two-tailed cluster-permutation, alpha p<.05, cluster alpha p<.05, N permutations=1000) (Maris & Oostenveld, 2007). The permutation distribution of t-values was constructed by storing the maximum summed absolute t-value at each iteration. Statistical significance of observed clusters was evaluated according to the p-value obtained by calculating the proportion of t-values under random permutation that were larger than the t-value of the observed cluster.

In addition to cluster-based, group-level permutation testing, specific hypotheses-driven comparisons between conditions in classifiers’ performance were additionally evaluated using paired-sample t-tests on AUC values averaged across specific time windows. This is especially warranted when quantifying relatively weak effects, because the latter statistical tests are more resilient to noise since the tests are not performed per sample, and furthermore, they can be more sensitive to short-lived effects that would otherwise not pass cluster thresholding (van Moorselaar & Slagter, 2019). Earlier work has identified two processing stages using the generalization across time approach: an early (<250-300ms) time-window, reflecting initial sensory encoding and a late processing stage (>300ms) associated with conscious report (e.g. Kaiser et al., 2016; Marti & Dehaene, 2017; Weaver et al., 2019). Based on this earlier work and based on observed time windows of significant decoding for letters and numbers in the localizer task of the current study (see the Result section), we focused our statistical analyses on two decoding clusters - one between 150-250ms and the other between 300-600ms. The diagonal AUC values within those two clusters were averaged separately and tested against each other using the paired-sample t-test. In cases where a specifically tested hypothesis did not indicate a significant result, using JASP software (JASP Team, 2020), we followed up that null-effect by a Bayesian equivalent of the same test in order to quantify the strength of evidence for the null hypothesis (H_0_) (Wagenmakers et al., 2018). By convention proposed by Jeffreys (1961), Bayes factors from 1 to 3 can be considered as anecdotal, 3 to 10 as substantial, and those above 10 as strong evidence in favor of H_0_.

Finally, we examined correspondence between our behavioral and decoding results. That is, we tested the extent to which the pattern of stimulus-specific target decoding (cross-task validation scheme) resembled behavioral results, reflecting conscious access across conditions. To that end, we used the same conditions that were entered into the behavioral analysis, but here, we used the average AUC decoding scores as the dependent measure in the repeated measures ANOVA. Again, one omnibus ANOVA was not possible since not all conditions had targets on the same TPs. We thus entered decoding scores into three repeated measures ANOVAs, investigating whether decoding scores across conditions reflect the AB, sparing, and blink reversal, respectively. We ran these three separate repeated measures ANOVAs, separately for early- and late-stage (150-250ms and 300-600ms) average AUC scores. Non-significant main and interaction effects were followed-up by a Bayesian equivalent of the same test in order to quantify the strength of evidence for the null hypothesis (H_0_) (Wagenmakers et al., 2018). Using JASP software (JASP Team, 2020), we conducted the Bayesian equivalent of the repeated measures ANOVA with the same within-subject factors as in the classical repeated measures ANOVA and computed exclusion Bayes factor (BF_excl_) across matched models, which indicates the extent to which data supports the exclusion of an interaction effect, taking all relevant models into account.

## 3 Results

### 3.1 Behavioral performance reveals flexibility of conscious access

We first aimed to replicate three key behavioral findings: the AB, sparing of conscious access, and AB reversal (Di Lollo et al., 2005; Olivers et al., 2007). Figure 1B shows percentages of correct target identification for our 8 conditions, which differed according to (1) the number of targets, and (2) their temporal position in the RSVP stream. In Figure 7, the behavioral results are also shown, but split up per conditions showing the AB (Fig. 7A), sparing (Fig. 7B), and attentional blink reversal (Fig. 7C). As can be seen in Figure 1B, participants identified single targets shown at the beginning and at the end of the stream equally well, suggesting that T1 performance was not significantly affected by target position in the stream alone. A paired-sample t-test revealed that there was no difference in performance for T1s presented on TP1 in condition 1 and T1s presented on late TPs in condition 8 (T_1_D_1_D_2_D_3_..T_2_: 87.6% vs. D_1_D_2_D_3_D_4_..T_2_: 87.9%, t_31_=−0.24, p=0.81, d=−0.043).

**Figure 1.**
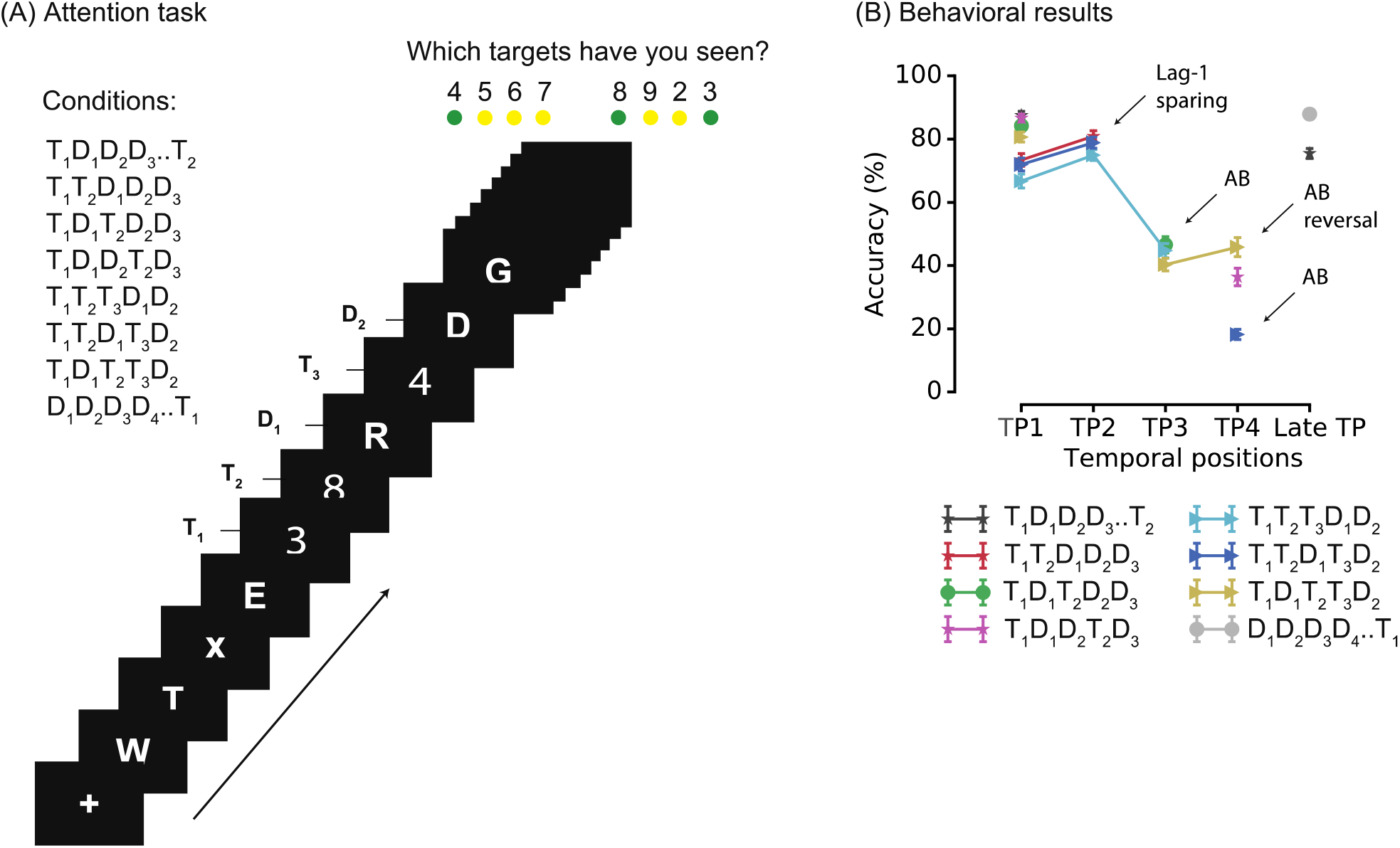
Attention task and behavioural results. (A) Conditions and the trial structure of the attention task. Each trial consisted of a sequence of rapidly presented letters in which 1-3 targets needed to be detected and reported at the end of the stream. Responses were registered using 8 marked keys on the keyboard, which spatially corresponded to 8 numbers shown on the computer screen in a specific order, for example 4 5 6 7 8 9 2 3, as shown in the figure. The order of stimuli on the response screen changed in every trial. (B) Percentage correct target identification for T1, T2 and T3 (given that T1 was correctly identified) as a function of temporal position and condition. Error bars represent SEM. As can be seen, our behavioral data demonstrate the presence of a robust AB, lag-1 sparing, AB reversal, but not of extended sparing (see also Figure 7).

As expected, we observed both a robust attentional blink to T2 and lag-1 sparing, as statistically captured by differences in T2 identification accuracy in 2-T conditions as a function of its temporal position with respect to T1 (main effect Temporal Position (2, 3, 4, and late (13-16)); F_3,93_=164.11, p<.001, η_p_^2^=0.841) and confirmed by planned follow-up pair-wise comparisons. Specifically, these revealed a clear AB to T2: T2s on TP3 (46.2%) in the T_1_D_1_**T_2_**D_2_D_3_ condition and on TP4 (35.7%) in the T_1_D_1_D_2_**T_2_**D_3_ condition were identified significantly less frequently than T2s at late TPs in the T_1_D_1_D_2_D_3_**T**_**2**_condition (75.3%) (t_31_=10.69, p=<.001, d=1.891; t_31_=13.14, p<.001, d=2.32). Furthermore, indicative of lag-1 sparing, we found that T2s on TP2 (lag-1) were detected more frequently than T2s shown late in the stream (T_1_**T_2_**D_1_D_2_D_3_: 80.7% vs. T_1_D_1_D_2_D_3_**T_2_**: 75.3%, t_31_=−2.82, p=.008, d=−0.49). Moreover, in line with some prior work (Dell’Acqua, Doro, Dux, & Losier, 2016; Di Lollo, Kawahara, Ghorashi, & Enns, 2005; Olivers, Hilkenmeier, & Scharlau, 2011), we found that T2 identification accuracy at lag-1 (TP2) was higher than T1 identification accuracy in the same condition (**T_1_T_2_**D_1_D_2_D_3_; 73.4% vs. 80.7%, t_31_ =-3.94, p<.001, d=−0.69).

We next examined whether sparing of conscious access extended beyond T2 to T3, as previous studies have demonstrated (Di Lollo et al., 2005; Olivers et al., 2007). A repeated measures ANOVA revealed that the pattern of results in the 3-T condition (T_1_T_2_T_3_D_1_D_2_) differed significantly from 2-T conditions (T_1_T_2_D_1_D_2_D_3_ and T_1_D_1_T_2_D_2_D_3_), as revealed by a Number of Targets (2-T vs. 3-T) x Temporal Position (1-3) interaction (F_2,62_=3.88, p=.026, η_p_^2^=0.11). A follow-up analysis showed that, in line with our earlier demonstration of target sparing on TP2 in the 2-T condition (T_1_T_2_D_1_D_2_D_3_), T2 accuracy was also spared on TP2 in the 3-T condition, and in fact, exceeded that of T1 (67%(T1) vs. 75%(T2), t_31_=−4.21, p<.001, d=-0.75). Nevertheless, our results suggested that the sparing did not extend to T3s presented immediately following T1 and T2. That is, a follow-up pair-wise comparison revealed that access to T3 on TP3 in the T_1_T_2_**T_3_**D_1_D_2_ condition was not significantly different from identification accuracy observed for targets on the same TP in the 2-T condition T_1_D_1_**T_2_**D_2_D_3_ (46.2% vs. 44.2%, t_31_=1.24, p=.225, d=0.22, BF_01_=2.64). These results thus suggest that conscious access was spared for the second, but not the third of three consecutive targets in the T_1_T_2_T_3_D_1_D_2_ condition in our study. This latter finding is unexpected given prior studies demonstrating extended sparing (Di Lollo et al., 2005; Olivers et al., 2007), and may be explained by the relative complexity of our target report procedure. Albeit speculative, having to remap which response button corresponded to which target number in each trial (necessary to decouple responses from target perception for our MVPA analyses) may have interfered with multiple target maintenance in working memory, and specifically affected T3 report.

Finally, we statistically verified the presence of attentional blink reversal (Olivers et al., 2007). As expected, T3s presented right after a T2 (T_1_D_1_T_2_**T_3_**D_2_) were detected more often compared to when they were separated by a distractor (T_1_T_2_D_1_**T_3_**D_2_) or compared to T2 at the same temporal position (T_1_D_1_D_2_**T_2_**D_3_), as indicated by a Number of Targets (2-T vs. 3-T) x Temporal Position (1, 3 and 4) interaction (F_2,62_=58.18, p <.001, η_p_^2^=0.65). This was confirmed by follow-up planned paired-sample t-tests which revealed that, although T2 identification was lower on TP3 in the 3-T condition than on the same TP in the 2-T condition (40.1% in T_1_D_1_**T_2_**T_3_D_2_ vs. 46.2% in T_1_D_1_**T_2_**D_2_D_3_, t_31_=4.68, p<.001, d=0.83), identification accuracy on TP4 in 3-T condition (T_1_D_1_T_2_**T_3_**D_2_, 44.8%) was significantly higher than accuracy on the same position in the 2-T condition (T_1_D_1_D_2_**T_2_**D_3_, 35.7%; t_31_=−5.92, p<.001, d=−1.05). Furthermore, T3 accuracy on TP4 was also higher than T2 accuracy on TP3 in the same 3-T condition (T_1_D_1_T_2_**T_3_**D_2_, 44.8% vs. T_1_D_1_**T_2_**T_3_D_2_, 40.1%; t_31_=-2.5, p=.019, d=-0.44). Lastly, we compared T3 accuracy on TP4 between two three target conditions which differed only in the temporal position of a preceding T2. Critically, when T3 immediately followed T2, as in the T_1_D_1_T_2_**T_3_**D_2_ condition, T3 accuracy was significantly higher compared to when T3 followed after T2 and a distractor, as in the T_1_T_2_D_1_**T_3_**D_2_ condition (44.8% vs. 18.1%; t_31_=−12.4; p<.001, d=-2.19). Together, these results reveal a clear reversal of the attentional blink.

Considered together, we replicated three key behavioral findings: the AB, sparing of conscious access for T2s presented immediately after T1 (i.e., lag-1 sparing), and AB reversal. However, we did not observe extended lag-2 sparing (to T3), possibly as noted above, due to our complex response protocol. The observed AB reversal for T3 in the T_1_D_1_T_2_T_3_D_2_ sequence in particular suggests that processes shaping conscious access are not necessarily temporally sluggish (e.g., determined by slow T1 encoding) (Marti & Dehaene, 2017; Marti et al., 2012; Sergent et al., 2005), but may depend on a fast information gating mechanism, e.g., dynamic excitation-inhibition feedback loops that modulate the strength of sensory representations as proposed by the boost and bounce theory (Olivers & Meteer, 2008; Olivers, et al., 2007). We next examined this hypothesis using EEG decoding analyses that allowed us to examine dynamic changes in the representational content of brain activity over time.

### 3.2 Decoding identity-specific target and distractor representations

Before examining neural representations of individual target and distractor stimuli, separately for T2 seen and unseen trials, and separately for boosted and bounced positions in the RSVP stream, we first verified that we could robustly decode individual letters and numbers using the localizer task data. As shown in Figure 2, individual numbers and letters could be decoded well above chance using classifiers trained on the localizer task data in the localizer task itself and, using cross-task classification, in the attention task. The resulting generalization across time matrices for the localizer task, shown separately for numbers and letters in Figure 2A, exhibited a mixture of diagonal and square shape decoding. Diagonal classification peaked at ~203 ms for numbers (AUC=53.09) and at ~156 ms (AUC=54.69) for letters. The decoding profile of stimulus-specific representations for numbers and letters also extended off diagonal after around ~450 ms, revealing the characteristic late-stage sustained squared-shaped profile (Carlson et al., 2013; King & Dehaene, 2014; Marti & Dehaene, 2017), which lasted for several hundred milliseconds, suggestive of stable stimulus representations across time. Note, however, that we did not observe early off-diagonal decoding, indicative of perceptual maintenance of early sensory representations (Marti & Dehaene, 2017; Meijs et al., 2019; Weaver et al., 2019).

**Figure 2.**
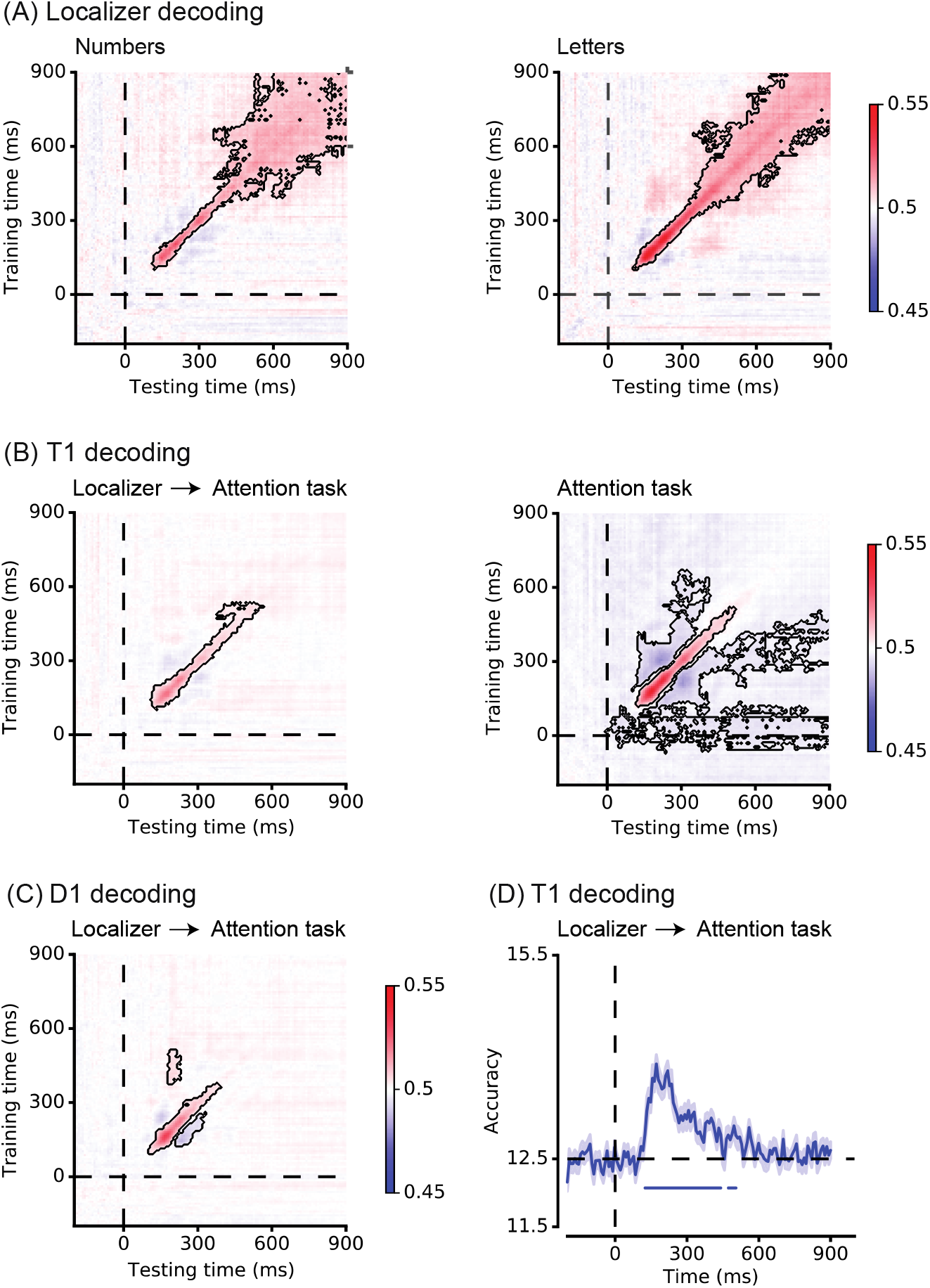
Time course of stimulus-identity decoding in the localizer (A, D) and attention task (B, C). (A) Generalization across time matrices based on within localizer task decoding reveal robust decoding of individual numbers and letters. Following a 10-fold cross-validation procedure, classifiers were trained on all time points and tested on all other time points, resulting in the generalization across time matrix for each stimulus category. The black contours on generalization across time matrices for number and letters indicate clusters of significant decoding of a stimulus identity (two-tailed cluster permutation test, alpha p<.05, cluster alpha p<.05, N-permutations=1000). (B) T1 identity decoding in the main attention task, based on training the classifiers on the localizer task data (left panel). T1 identity decoding based on a 10-fold cross-validation scheme, using the attention task data (right panel). (C) D1 identity decoding in the main attention task, based on localizer task classifier (cross-task validation procedure). (D) Diagonal T1 identity decoding in the attention task based on the localizer task classifier, using accuracy to evaluate classification performance. Note that classification accuracy and AUC scores, which are used as classification metric throughout the paper, show highly similar decoding pattern (see Fig. 3A) All plots show classification performance averaged over all conditions and all participants. Time 0 ms in all plots corresponds to T1 presentation time, except for data shown in panel C and the D1 diagonal line in panel D, where D1 time course was shifted back one temporal position for visualization purposes.

Cross-task classification (i.e., localizer to attention task data classification) also showed robust decoding of both target (number) and distractor (letter) stimulus identity in the attention task (Fig. 2B-C), with representations of successive stimuli partially overlapping in time (Fig. 3). T1-specific patterns of activity emerged around 116 ms, with diagonal classifier performance peaking at ~172 ms (Fig 2B, left panel). A similar decoding profile was observed for D1s: identity specific patterns emerged around 100 ms, with diagonal decoding peaking at ~164 ms. The cluster of significant T1 decoding was notably more temporally extended in comparison to D1 decoding, lasting until 528 ms versus 378 ms, respectively, likely reflecting stimulus differences in task relevance (i.e., report numbers). Therefore, and in line with previous work that identified two similar processing stages using MVPA analyses (e.g. Marti & Dehaene, 2017; Meijs et al., 2019; Weaver et al., 2019) (see Fig. 2A-B), in subsequent statistical analyses comparing decoding accuracy in different conditions, we averaged diagonal AUC values across two time windows that capture these two processing stages: an early 150-250 ms time-window, reflecting initial sensory encoding, and a later 300-600 ms time-window, associated with conscious report.

**Figure 3.**
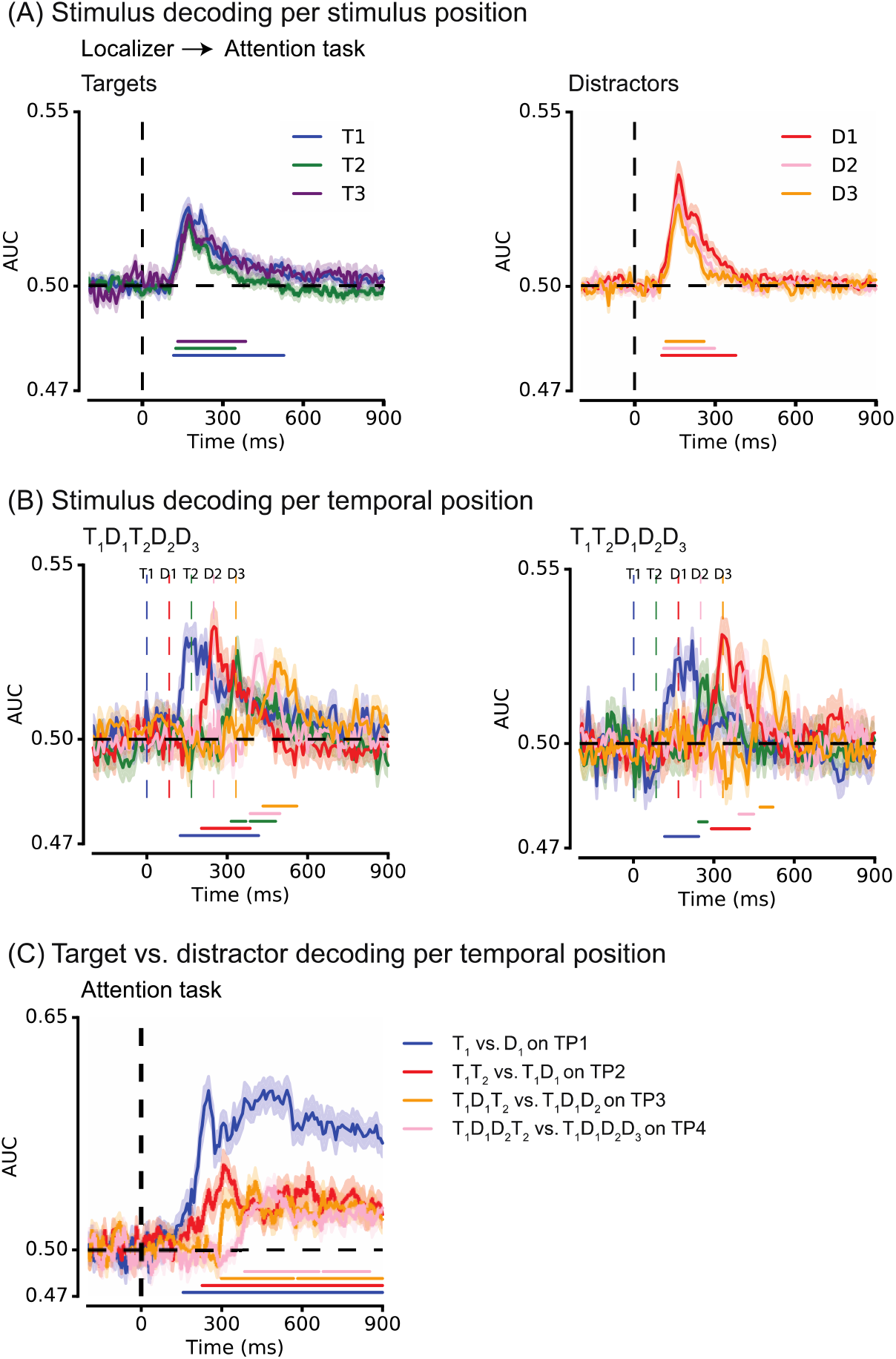
Time-resolved identity decoding as a function of stimulus class (target, distractor) and temporal position in the attention task. (A) Target and distractor identity decoding in the attention task, based on the localizer task classifier (cross-task validation), as a function of the target or distractor number in the RSVP stream of the attention task. EEG data of all stimuli except T1 were locked to the presentation time of a given stimulus and then shifted to T1 presentation time, given the variable presentation times of stimuli across conditions. (B) Cross-task decoding for target and distractor stimuli as a function of temporal position for two conditions. The colored dashed vertical lines indicate objective presentation times of each stimulus in a given condition. Note that because letter identity was better decodable than number identity in the localizer task, target and distractor identity decoding in the attention task cannot be directly compared. This figure simply demonstrates the ability of our approach to decode each individual stimulus in the RSVP stream. (C) Binary target versus distractor diagonal decoding per temporal position using 10-fold validation scheme. At each temporal position (TP), target and distractor labels were obtained from 2 conditions: TP1 – **D_1_**D_2_D_3_D_4_..T_1_ vs. **T_1_**D_1_D_2_D_3_..T_2_; TP2 – T_1_**D_1_**D_2_D_3_..T_2_ vs. T_1_**T_2_**D_1_D_2_D_3_; TP3 – T_1_D_1_**D_2_**D_3_..T_2_ vs. T_1_D_1_**T_2_**D_2_D_3_; TP4 – T_1_D_1_D_2_**D_3_**..T_2_ and T_1_D_1_D_2_**T_2_**D_3_. In all plots, the colored horizontal lines indicate periods of significant decoding with respect to chance (two-tailed cluster permutation test, alpha p<.05, cluster alpha p<.05, N-permutations=1000). All plots show classification performance averaged over all participants.

It should be noted that overall, early decoding accuracy for letters (distractors) was higher than for numbers (targets) in both the localizer task (see early diagonal decoding scores for numbers and letters in the localizer task in Fig. 2A) and the attention task (Fig 3A). As in the localizer task, both letters and numbers were decoded as targets, these differences in early decoding accuracy cannot reflect differences in task relevance, and likely reflect the fact that letters and numbers are processed in different brain regions (Carreiras, Quiñones, Hernández-Cabrera, & Duñabeitia, 2015), whose activity may be differentially measurable on the scalp (e.g., due to anatomical differences in how they are oriented with respect to the scalp). For this reason, target and distractor decoding is not statistically compared directly in any of the reported analyses in the further text.

To summarize, we could robustly decode, in parallel (see Fig. 3B), individual numbers and letters in the attention task and replicate previous reports of two distinct processing stages (Kaiser et al., 2016; Marti & Dehaene, 2017; Meijs et al., 2019). We next examined 1) if, how and when in time (early vs. late) these sensory representations differed between T2 seen and unseen trials and 2) if they were modulated depending on whether the stimuli were presented on boosted or bounced positions in the RSVP stream (i.e. depending on the category of the preceding stimulus: target or distractor).

### 3.3 Early identity-specific stimuli representations are not ‘boosted’ or ‘bounced’

In contrast to limited-capacity theories that propose that the attentional blink to T2 is caused by late-stage T1 encoding (Lagroix et al., 2012; Sergent et al., 2005), the boost and bounce theory posits that the attentional blink is due to D1-related dysfunctional gating of information, and hence the theory predicts differences in the neural representation of D1 in T2 seen vs. unseen trials (Olivers & Meeter, 2008). Therefore, we next examined possible differences in early and late sensory representation of T1, D1, and T2 as a function of whether T2 was seen or not. By splitting the analysis for T2 seen and unseen trials, we aimed to test 1) whether the duration and/or the strength of T1 processing differs between T2 seen and unseen trials as limited-capacity accounts would predict (i.e. resulting in longer and/or stronger T1 representations in T2 unseen trials), 2) whether, as proposed by the boost and bounce account, early D1 representations are amplified in T2 unseen versus seen trials and 3) if T2 representations are weaker when T2 is not seen vs. seen, as both accounts would predict. To foreshadow our results, shown in Figure 4, these analyses yielded an unexpected link between the strength of stimulus-specific representations and conscious access. First, we found that T1 stimulus representations were significantly stronger on T2 seen versus unseen trials both during the early (t_31_=2.78, p=.01) and late (t_31_=2.62, p=.01) time window. Further, contrary to what the limited capacity account would predict, we also found that T1 stimulus-specific representations could be decoded for a longer period of time on T2 seen trials than on T2 unseen trials. In both trial categories, T1 identity could be decoded above chance from ~117 ms onwards, but T1 decoding was significant until ~433 ms in T2 seen versus ~275 ms in T2 unseen condition (see Figure 4A), although the magnitude of this difference was not significant.

**Figure 4.**
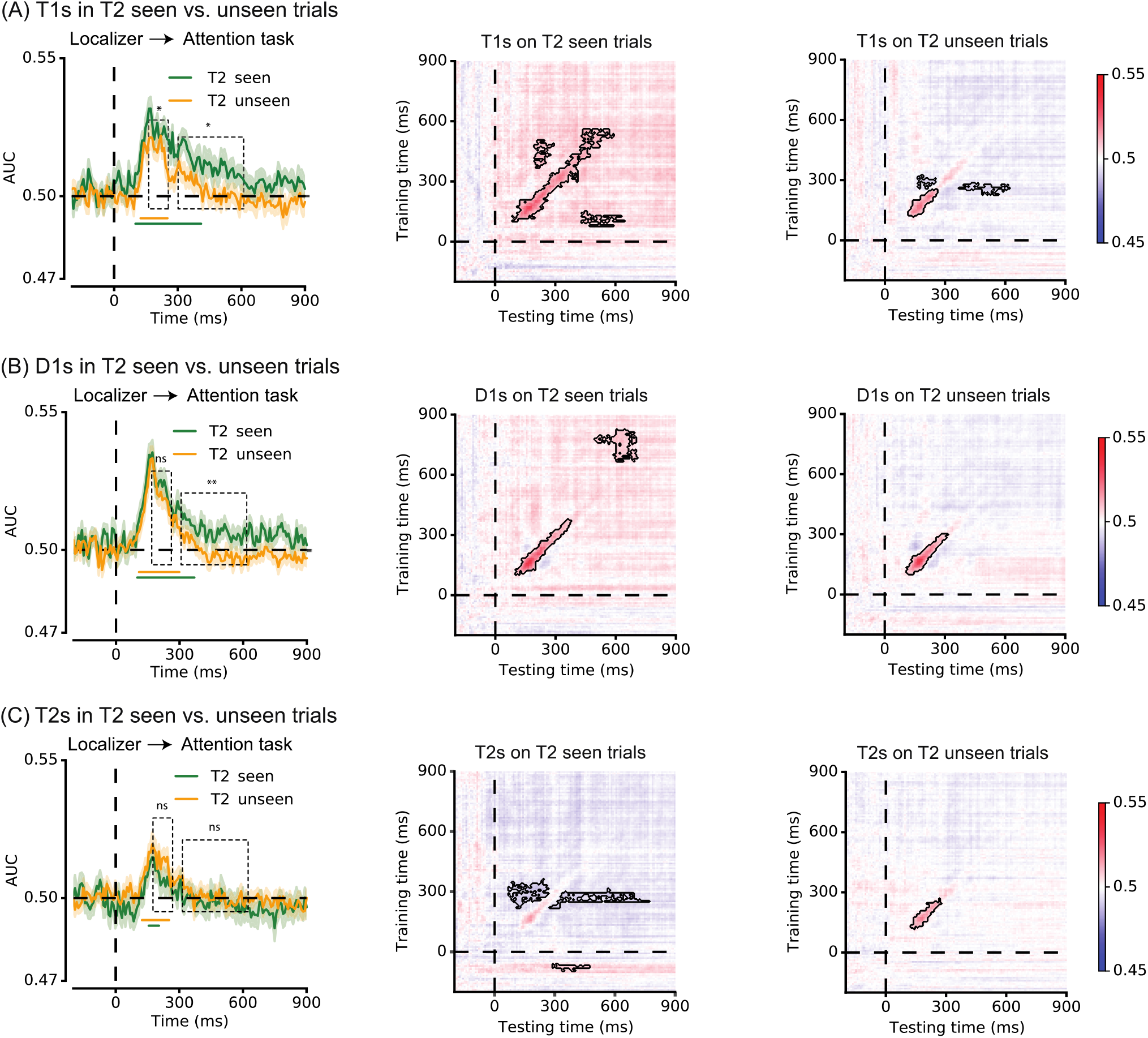
Time-resolved decoding of T1, D1 and T2 stimuli, separately for T2 seen and unseen trials. Cross-task diagonal decoding and generalization across time of stimulus identity for T1 (A), D1 (B) and T2 (C) in T2 seen and unseen trials based on T1D1T2D2D3, T1D1D2T2D3 and T1D1T2T3D2 conditions. In all diagonal decoding plots, the colored horizontal lines indicate periods of significant decoding with respect to chance (two-tailed cluster permutation test, alpha p<.05, cluster alpha p<.05, N-permutations=1000). The black dashed rectangles indicate the early and late-stage time windows used to compare AUC decoding scores between conditions. All plots show classification performance averaged over all participants. This figure shows that conscious access to T2 was associated with stronger early and late stage T1 representations and stronger late stage D1 representations, but no differences in T2 neural representation itself. In all figure panels, time 0 ms corresponds to the presentation time of the stimulus of interest (e.g., T1 onset latency in A).

Next, we tested differences in D1 representation between T2 seen and unseen trials. While early D1 representations did not significantly differ between T2 seen and unseen trials (t_31_=1.00, p=.32, BF_01_=3.34), the strength of D1 representations, like T1 representations, was significantly higher in trials in which T2 was seen vs. unseen during the late (300-600 ms) processing stage (t_31_=2.83, p<.01). Additionally, a group-level cluster-based permutation test indicated that diagonal D1 decoding was more extended in time on T2 seen versus unseen trials (lasting until ~560 ms vs. ~299 ms). Thus, we found that both T1 and D1 were better decodable in trials in which T2 was seen vs. blinked. These results are unexpected from a limited capacity perspective, which assumes stronger or longer-lasting late-stage processing (representation) of T1 in T2 unseen, rather than T2 seen, trials, but also from the boost and bounce account, which would propose that the AB is associated with attentional selection of D1 in T2 unseen trials. Yet, our results suggest that both T1 and D1 were more strongly represented on T2 seen versus unseen trials.

Furthermore, we found that early T2 representations did not differ in strength as a function of whether T2s were seen or unseen. Figure 4C shows classifiers’ AUC scores for T2 decoding in three conditions, T_1_D_1_**T_2_**D_2_D_3_, T_1_D_1_D_2_**T_2_**D_3_ and T_1_D_1_**T_2_**T_3_D_2_. The reason for collapsing across these three conditions was that in each of these conditions, T2 followed T1 after one or two distractors and individual conditions had too low trial numbers to achieve robust identity decoding. At the behavioral level, T2 accuracy in each of these conditions was very comparable (Figure 1B). Both cross-task and within attention task T2 decoding did not provide evidence for differences in T2 seen and unseen AUC scores during the early 150–250 ms time window (cross-task: t_31_=−1.71, p =.098, BF_01_=1.45; within-task: t_30_=−1.12, p=0.28, BF_01_=2.95) or late 300–600 ms time-window (cross-task: t_31_=−0.44, p=.66, BF_01_=4.84; within-task: t_30_=0.98, p=0.34, BF_01_=3.37) (Figure 4C; within-task decoding is not shown in the figure). Note that accuracy of late-stage T2 decoding, both on and off diagonal, was close to chance. This weak late-stage decoding likely reflects the fact that employed classifiers were tuned to identity-specific patterns of activation, and thus less sensitive to later processes associated with encoding and conscious access. It is conceivable that in a context with multiple targets, representational codes of later targets become more variable in latency or in format in which a target is encoded, which would thus render robust classification between the tasks difficult. Overlap from preceding items may have also interfered with T2 decoding.

We did uncover differences between T2 seen and unseen processing using two different analysis approaches. First, replicating prior work (Sergent et al., 2005; Sigman & Dehaene, 2008; Vogel et al., 1998), we found that the magnitude of the T2-evoked centro-parietal P3 ERP component was significantly larger on T2 seen compared to unseen trials (600-800ms post-T1: t_31_=5.26, p<.001) (Figure 5A). As can been seen in this figure, the T2-evoked P3 was preceded in time by the T1-evoked P3 around 300-550 ms post-T1. That this reflects T1-evoked activity is supported by the fact that this first positivity was also observed in long lag trials (T_1_D_1_D_2_D_3_..T_2_), in which T2 was presented much later, and in which hence, as expected, no second positivity was observed between 600 and 800ms post-T1. The T1-evoked P3 did not differ between T2-seen and unseen short-lag trials (t_31_=0.81, p=0.43).

**Figure 5.**
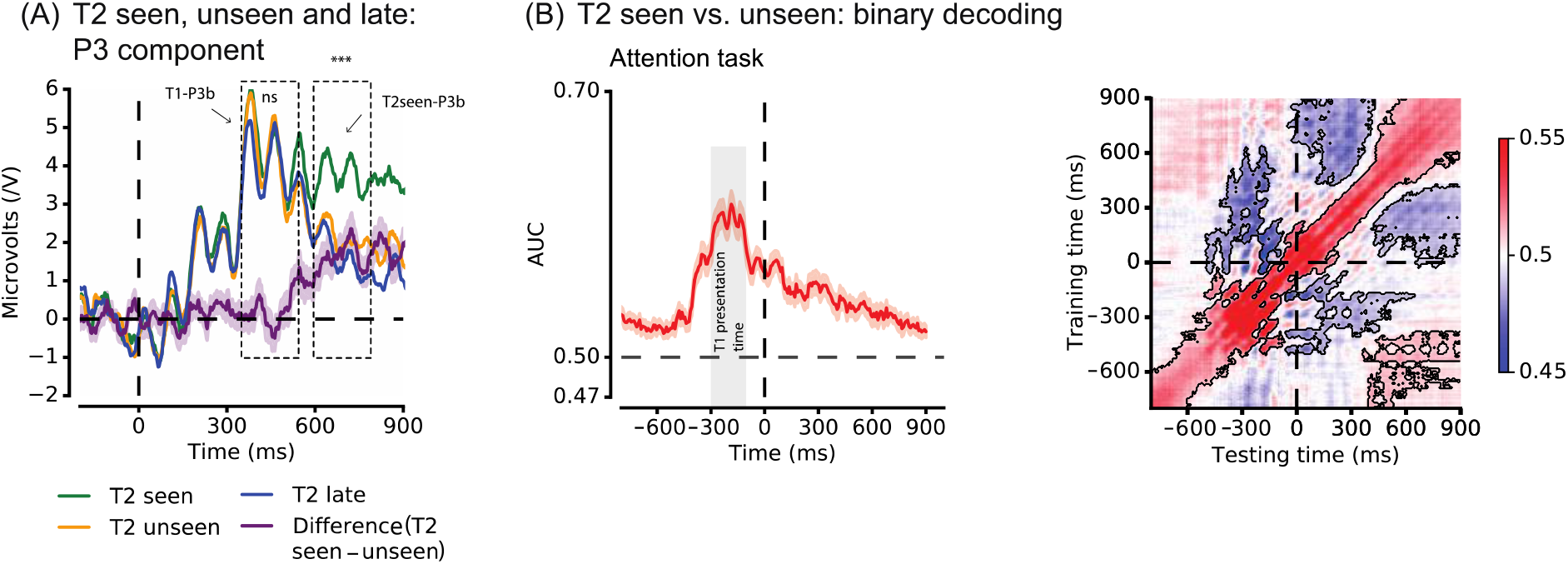
ERP analysis and time-resolved binary decoding for T2 seen and unseen trials. (A) Seen T2s evoked a larger P3b than unseen T2s. Shown is the centro-parietal P3 component measured on channels POz, Pz, CPz, CP1, CP2, P1, P2, PO3 and PO4, separately for T2 seen and unseen trials in the T_1_D_1_T_2_D_2_D_3_ condition (green and orange lines), and for the T_1_D_1_D_2_D_3…_T_2_ condition in which T2 appears on a late temporal position (blue line), given that T1 was correctly identified. The purple line is the difference waveform between T2 seen and T2 unseen waveforms in the T_1_D_1_T_2_D_2_D_3_ condition. Time 0 ms corresponds to T1 presentation time. (B) T2 seen versus unseen decoding based on the main attention task data, using three conditions (T_1_D_1_T_2_D_2_D_3_, T_1_D_1_D_2_T_2_D_3_ and T_1_D_1_T_2_T_3_D_2_). In this analysis, using a 10-fold cross validation procedure, a classifier was trained to distinguish between two classes of labels: T2 seen versus unseen (T2 identity was therefore irrelevant). Seen T2s were differently represented than unseen T2s for up to 900ms post-T2 presentation, and conscious access to T2 was also associated with differences in the pattern of brain activity prior to T2 presentation. Time 0 ms corresponds to T2 presentation time.

Second, classifiers trained to decode whether a T2 was seen or unseen in the main attention task, irrespective of the T2 identity (classifier labels: T2 seen vs. T2 missed; i.e., T2 identity was irrelevant) revealed clusters of significant decoding scores for over 900ms after T2 presentation (Fig. 5B), confirming that the neural signal contained information related to conscious T2 access throughout the trial. Interestingly, this analysis also showed enhanced decoding well before T2 presentation suggesting that, besides T2 processing, neural activity prior to T2 presentation also predicted whether T2 would be seen or not. Diagonal decoding started rising above chance approximately around the onset of the first item in the RSVP stream (between ~580 ms and ~500 ms) and reached a maximum around T1 presentation time (approximately −300 to −100 ms before T2 presentation time; Figure 5B). This finding may corroborate previous findings suggesting that baseline fluctuations in neural excitability and attention across trials shapes the likelihood of conscious access to a significant extent (Iemi, Chaumon, Crouzet, & Busch, 2017; Mathewson, Gratton, Fabiani, Beck, & Ro, 2009). It could also reflect differences in temporal expectation of T1 (which had a fixed position in the stream) between blink and no-blink trials.

To summarize, we found that the attentional blink was associated with weaker representations of T1 and D1, rather than enhanced or prolonged late-stage T1 encoding, as limited capacity accounts propose, or amplified D1 representations, as the boost and bounce theory assumes. Rather, we could better decode T1 both during early- and late-stage processing and D1 during late stage processing in trials in which T2 was *seen*.

Next to examining T1 and D1 stimuli representations as a function of T2 visibility, we investigated whether target and distractor representations are modulated according to whether the position at which they are presented in the RSVP stream is boosted (following a target) or bounced (following a distractor that is itself boosted). As noted above, limited-capacity accounts have trouble explaining the behavioral observations of extended sparing and AB reversal, which the boost and bounce theory links to dynamic changes in top-down attentional modulation. According to the latter account, a rapidly responding gating system enhances target and suppresses distractor representations, but when these stimuli are quickly followed in time by another stimulus, this also affects their early representation and thereby their reportability. To investigate if the sensory representation of a stimulus is affected by the nature of the preceding stimulus (target or distractor), we first focused on the phenomenon of AB reversal - enhanced identification of T3s when directly preceded by a target (T_1_D_1_T_2_**T_3_**D_2_) vs. a boosted distractor (T_1_T_2_D_1_**T_3_**D_2_). The boost and bounce account predicts that the sensory representation of T3 should be enhanced when directly preceded by T2 compared to when it is preceded by a distractor. However, although T3 decoding was numerically stronger along the diagonal in the T_1_D_1_T_2_**T_3_**D_2_ versus T_1_T_2_D_1_**T_3_**D_2_ condition during late-stage processing, the difference between conditions did not reach significance in the early or late time window (early stage: t_31_=0.31, p=.76, BF_01_=5.06; late stage: t_31_=1.44, p=.16, BF_01_=2.09) (Figure 6A).

**Figure 6.**
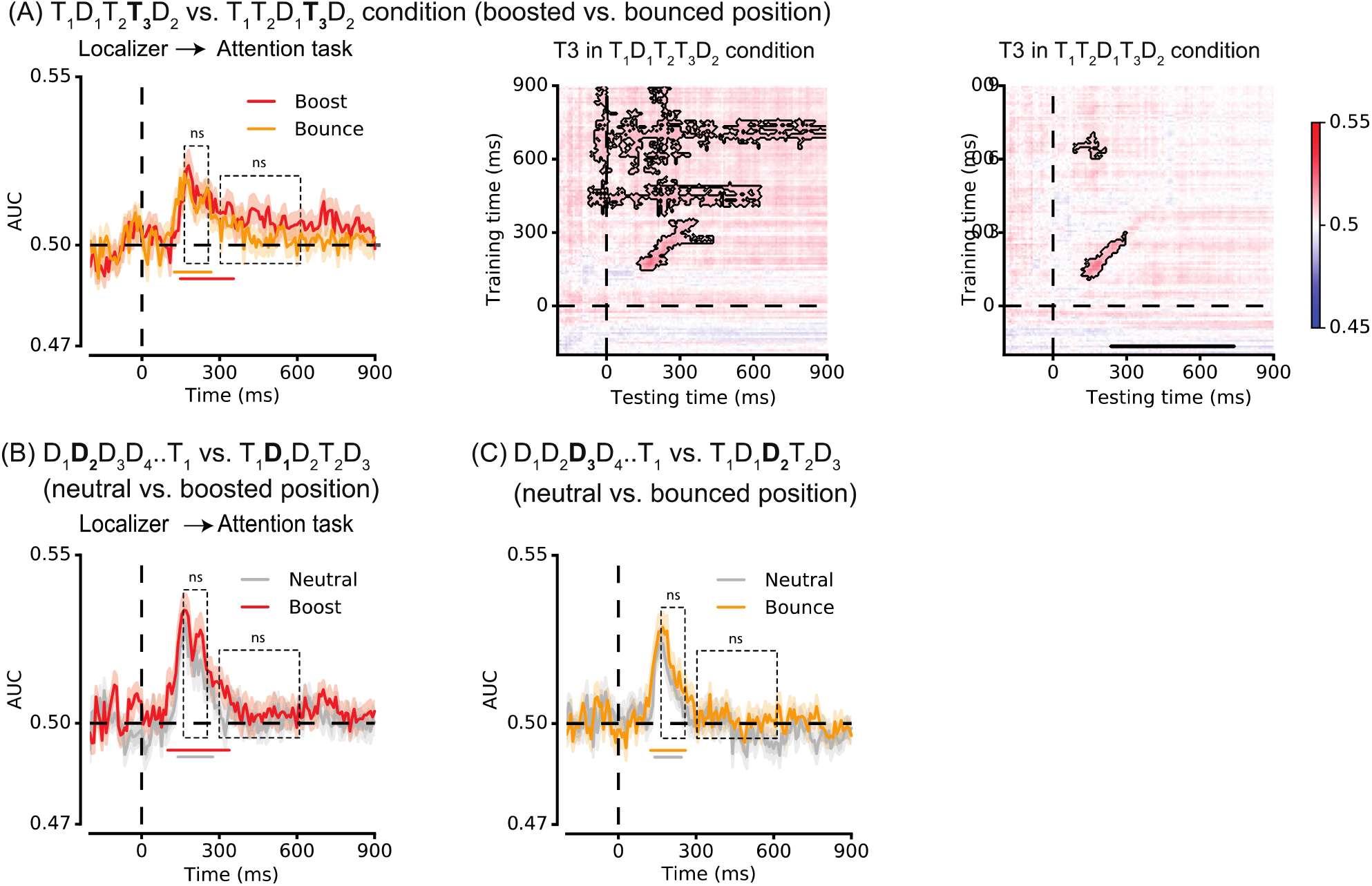
Time-resolved stimulus identity decoding does not reveal differences in representational strength on ‘boosted’ versus ‘bounced’ positions. (A) Cross-task diagonal T3 identity decoding in condition T_1_D_1_T_2_**T_3_**D_2_ (T3 on boosted position) versus T_1_T_2_D_1_**T_3_**D_2_ (T3 on bounced position), and the generalization across time matrix of T3 identity decoding for each condition separately. Cross-task diagonal identity decoding for distractors presented (B) on temporal position 2 (TP2) in the D_1_**D_2_**D_3_D_4_..T_1_ condition (neutral position) versus in the T1**D1**D2T2D3 condition (boosted position), and (C) on temporal position 3 (TP3) in the D_1_D_2_**D_3_**D_4_..T_1_ condition (neutral position) versus the T_1_D_1_**D_2_**T_2_D_3_ condition (bounced position). In all plots, the colored horizontal lines indicate periods of significant decoding with respect to chance (two-tailed cluster permutation test, alpha p<.05, cluster alpha p<.05, N-permutations=1000). The black dashed rectangles indicate time periods used for statistical comparisons between conditions. All plots show classification performance averaged over all participants. In all figure panels, time 0 ms corresponds to the presentation time of the stimulus of interest (e.g., T3 onset latency in A).

Lastly, we investigated whether distractor representations may be modulated depending on whether they were preceded by a target (T1) and/or a distractor (Figure 6B & C). Specifically, we compared distractor representations in the T_1_D_1_D_2_T_2_D_3_ condition and D_1_D_2_D_3_D_4_..T_1_ condition, separately for distractors shown on TP2 and TP3. Note that both positions in the D_1_D_2_D_3_D_4_..T_1_ condition can be considered neutral since TP2 and TP3 distractors were always preceded by other distractors. In the T_1_D_1_D_2_T_2_D_3_ condition, distractors on TP2 directly followed T1 (i.e., ‘boosted’ D1s), while those on TP3 directly followed D1 (i.e., ‘bounced’ D2s). Statistical comparison of decoding scores for distractor representations on boosted and neutral TP2 positions suggested that those did not differ significantly during the early (t_31_=−1.57, p=.13, BF_01_=1.75) and late (t_31_=-0.67, p=.51, BF_01_=4.3) time interval. The same was true for distractors on neutral and bounced positions. That is, the chance to decode distractor representations was not statistically different on neutral versus bounced TP3 positions during the early (t_39_=-1.33, p=.19, BF_01_=2.37) or late (t_39_=-0.68, p=.50, BF_01_=4.27) time period of decoding. Taken together, our results provide no convincing evidence for a modulation of identity-specific representations as a function of the task-relevance of the preceding stimulus.

### 3.4 Target decoding does not resemble target report accuracy across conditions

Post-hoc trial sorting and analysis based on an outcome measure (seen vs. unseen), as was done in the above decoding analyses, can create confounds in condition comparisons (Shanks, 2017), and moreover, reduces the number of trials per analysis cell. Moreover, above, we contrasted decoding accuracy between two specific conditions (e.g., D1 decoding in T2 seen vs. unseen trials), while there is naturally more information in result patterns across multiple conditions. We therefore next directly evaluated whether the pattern of decoding results exhibits three key events, namely the AB, lag-1 sparing and AB reversal, which we observed behaviorally. To that end, we ran three separate repeated measures ANOVAs, identical to those we ran on behavioral data, separately for early- and late-stage (150-250 ms and 300-600 ms) average AUC scores. In this way, we could determine whether the strength of (early or late stage) target decoding resembles target identification accuracy, taking multiple conditions into account, as we did previously for the behavioral analysis. Figure 7 displays the patterns of target identification accuracy and early and late target decoding accuracy separately for the conditions used to identify the AB, lag-1 sparing and AB reversal.

**Figure 7.**
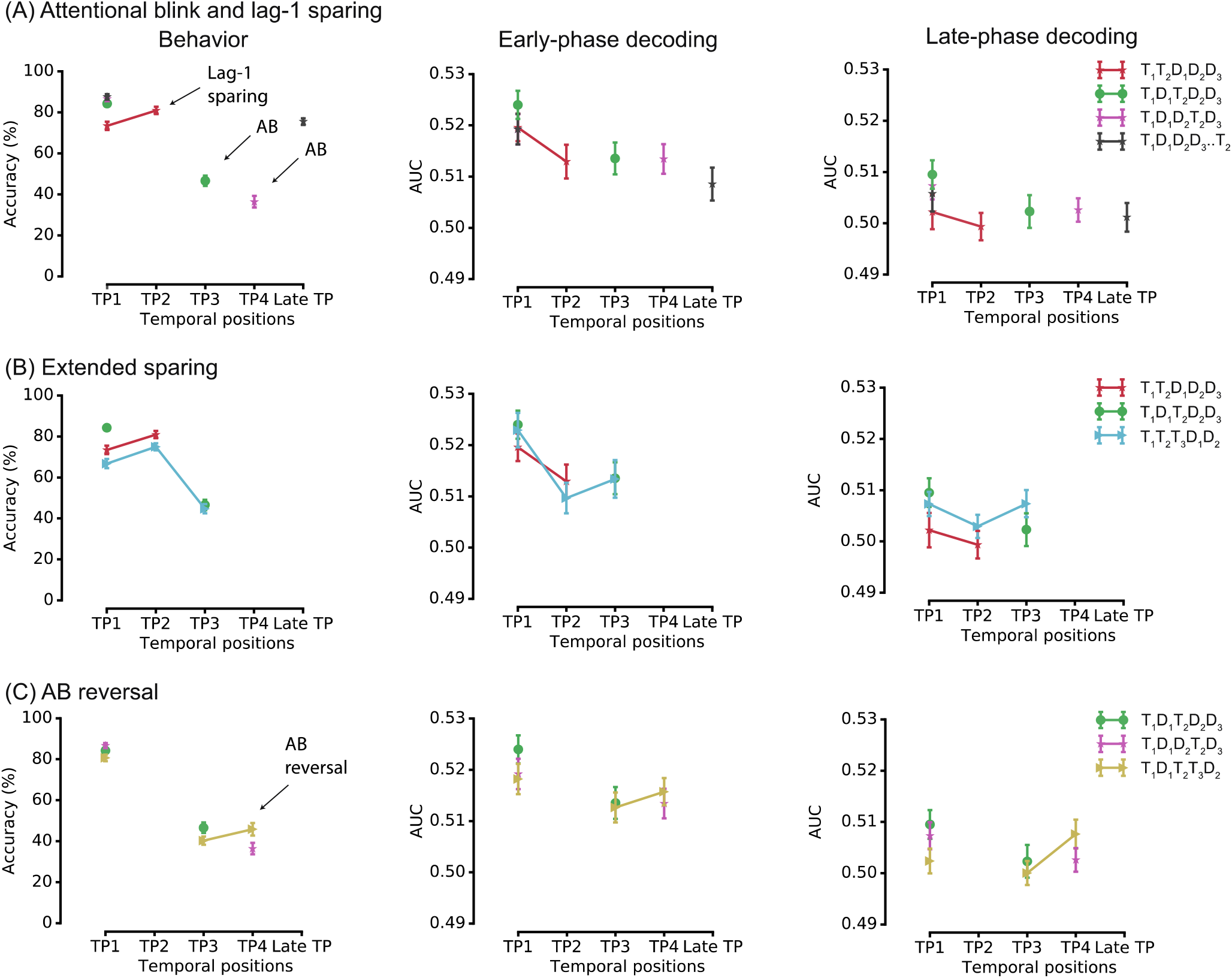
Subjects’ behavioral and classifiers’ decoding performance in identifying targets as a function of temporal position (TP). (A) Average behavioral accuracy (left column) and AUC decoding scores, separately for early-phase (middle column) and late-phase (right column) decoding, shown for conditions that at the behavioral level demonstrate the presence of an AB and lag-1 sparing in the attention task, (B) conditions that at the behavioral level demonstrate the presence of lag-1 sparing, but not of extended sparing in the attention task, (C) and conditions that at the behavioral level demonstrate AB reversal.

First, we tested whether early stage T2 decoding varied across the 2-T conditions, reflecting the behavioral pattern of the AB. A one-way repeated measures ANOVA on the early decoding data (150-200 ms), showed that early T2 decoding accuracies did not differ across temporal positions (F_3,93_=0.49, p=.69, BF_01_=12.56). This was also the case for the late-stage (300–600 ms) decoding scores, as reflected by the absence of a main effect of Temporal Position in a one-way repeated measures ANOVA on AUC scores (F_3,93_=0.38, p=.77, BF_01_=15.06). Thus, the AB was not reflected in early or late T2 decoding scores (Figure 7A). Next, we tested whether the pattern of target decoding in 2-T and 3-T conditions (T_1_T_2_D_1_D_2_D_3_, T_1_D_1_T_2_D_2_D_3_, T_1_T_2_T_3_D_1_D_2_) exhibited sparing for targets, as observed behaviorally for T2s. A two-way repeated measures ANOVA using AUC decoding scores as the dependent variable did not provide evidence for sparing (higher decoding scores of targets preceded by another target vs. distractor), as decoding scores did not differ significantly between 2-T and 3-T conditions across temporal positions, as indicated by a non-significant Number of Targets x Temporal Position interaction for early-stage decoding scores (F_2,62_=0.26, p=.77, BF_01_=8.20) and late-stage decoding scores (F_62,2_=0.11, p=.89, BF_01_=9.88) (Figure 7B). The main factor Number of Targets was also not significant for early- or late-decoding scores (early: F_1,31_=8.99e-4, p=.98, BF_01_=6.51; late: F_1,31_=3.85, p=.06, BF_01_=1.01) suggesting that there was no difference in decoding between conditions with two versus three targets. Yet, decoding was found to be significantly lower for later temporal positions, as indicated by the main effect of Temporal Position for early decoding scores (F_2,62_=4.61, p=.01), but not for late decoding scores (F_2,62_=0.97, p=.39, BF_01_=6.14) (Figure 7B). Finally, we also did not find convincing evidence for AB reversal using target decoding scores (Figure 7C). A two-way repeated measures ANOVA again revealed that the Number of Targets x Temporal Position interaction was not significant for the early decoding (F_2,62_=1.26, p=.29, BF_01_=3.85), although the interaction was trend-level significant for the late decoding scores (F_2,62_=2.73, p =.07), supported by weak evidence for the null hypothesis as revealed by the Bayes factor of BF_01_=0.75. Suggested by the null-effect of the factor Number of Targets for early decoding (F_1,31_=0.54, p =.47, BF_01_=5.1) and late decoding (F_1,31_=0.78, p=.39, BF_01_=4.82), decoding was not modulated by the number of targets in analyzed conditions, but decoding was affected by the temporal position of targets, although only during early-stage decoding (main effect of Temporal Position, early: F_1,31_=4.82, p =.01, late: F_1,31_=1.37, p =.26, BF_01_=4.43).

In summary, we found that target representational strength did not reliably reflect the attentional blink, sparing, or AB reversal as differences in decoding across corresponding conditions were statistically not significant.

## 4 Discussion

The present study aimed to enhance understanding on how attentional selection shapes conscious access under conditions of rapidly changing input. Using an attention task and multivariate decoding of individual target- and distractor-defining features, we specifically examined dynamic changes in the representation of targets and distractors as a function of conscious T2 access and the task-relevance (target or distractor) of the preceding item in the RSVP stream. At the behavioral level, we found compelling evidence for a flexible gating mechanism, replicating previous findings (Di Lollo et al., 2005; Lunau & Olivers, 2010; Olivers et al., 2011, 2007). That is, we found a significant impairment in conscious access to targets that were preceded by one or two distractors (i.e., the AB), but striking facilitation of conscious access to targets shown directly after another target (i.e., lag-1 sparing and AB reversal). Yet, our neural data did not provide convincing evidence for selection-related feedback effects on early-stage visual representations as a determinant of conscious access under rapidly changing input conditions (Olivers & Meeter, 2008): early-stage representations of D1 did not differ between trials in which T2 was seen versus blinked, nor was the early-stage representation of T3 affected by whether T3 was preceded by another target or a distractor. Furthermore, overall, the strength of early stimulus representations across conditions exhibited little variability, and our statistical models suggested that the general pattern of early, as well as late, decoding results did not resemble that which we observed in behavioral performance. Our findings thus indicate that conscious access under rapidly changing input conditions may be dependent on other mechanisms than rapid top-down modulation of early low-level sensory representations. Notably, conscious access to T2 was associated with stronger early- and late-stage T1 representations, as well as stronger late-stage D1 representation, suggesting that both differences in T1 and D1 processing may precede the attentional blink to T2. These findings have implications for theories of the attentional blink and consciousness more generally, discussed below.

Our findings corroborate previous work showing that multiple sensory representations can coexist in patterns of neural activity for a few hundred milliseconds, presumably at different (early) stages of processing (Grootswagers et al., 2019; Marti & Dehaene, 2017; Tang et al., 2020). Temporal decoding profiles of target and distractor stimuli were robust and remarkably similar up to approximately 250 ms, confirming that early stages of visual processing are common to all stimuli - seen or unseen - entering the visual system, while late-stage processing is selective to consciously perceived stimuli (Marti & Dehaene, 2017). One of our main findings was that conscious access to T2 was associated with stronger early- and late-stage T1 representations, as well as stronger late-stage D1 representation, indicating that encoding of both T1 and D1 may dynamically affect conscious perception and access of subsequent stimuli (Fahrenfort et al., 2017; Martin, Cox, Scholl, & Riesenhuber, 2019). A striking aspect of our findings was the direction of the observed differences, namely, conscious access to T2 was associated with enhanced T1 and late-stage D1 representations. Thus, when T2 was blinked, the strength of early and late-stage T1 representations and of late-stage D1 representations was lower. This observation is not only difficult to reconcile with theories that postulate that enhanced T1 encoding causes the AB and that do not assign a critical role to D1 in the AB (e.g. Chun & Potter, 1995), but are also surprising in light of accounts that posit that the attentional blink is due to D1 accidentally being boosted into working memory (Olivers & Meeter, 2008) and thus would predict D1 processing to be enhanced or prolonged in T2 blink trials, not in T2 seen trials, contrary to what we observed.

One explanation for our findings, which could reconcile them with the larger literature, is that an enhanced sensory representation may reduce the time necessary for higher-level encoding of a stimulus into a durable format, and thus indicates more efficient working memory encoding. The serial token/simultaneous type model (Bowman, Wyble, Chennu, & Craston, 2008) actually makes this prediction. That is, this model proposes that a reciprocal relationship between T1 bottom-up trace (or stimulus) strength and encoding time underlies the AB. Specifically, stronger T1 representations necessitate less attentional enhancement, so that attention can be more quickly reallocated to T2, rendering it more likely that T2 will be perceived. The serial token/simultaneous type model would hence predict an initial stronger T1 representation in no-blink trials, as we find here. In the boost and bounce model (Olivers & Meeter, 2008), a more robust bottom-up T1 representation could also reduce the need for top-down amplification due to stronger initial evidence for its presence, which would consequently also reduce the strength of the subsequent D1-evoked bounce response. However, our results do not show any evidence for distractor-evoked suppression of the representation of following items.

If an enhanced bottom-up sensory representation of T1 reduced the time necessary for higher-level encoding of T1 into a durable format, one may also expect the T1-evoked P3b to peak earlier or be smaller in amplitude in no-blink compared to blink trials. Yet, the T1-evoked P3b did not differ between T2 seen and unseen trials in the present study. While some ERP studies have reported a larger T1-evoked P3b in T2 blink trials (e.g., Kranczioch, Debener, Maye, & Engel, 2007; Martens, Elmallah, London, & Johnson, 2006; Shapiro, Schmitz, Martens, Hommel, & Schnitzler, 2006), other ERP studies reported a delayed T1-evoked P3b (e.g., Martens, Munneke, et al., 2006; Sergent et al., 2005). As in the current study, yet other studies did not observe any difference in the amplitude or latency of the T1-evoked P3b between blink and no-blink trials (e.g., Craston, Wyble, Chennu, & Bowman, 2009; Kihara & Kawahara, 2008; Slagter et al., 2007, pre-retreat data). Thus, AB-related differences in late-stage T1 processing are not consistently observed across studies. Notably, a novel line of evidence suggests that the P3b component does not track perception and encoding, but rather post-perceptual processes (e.g., decision making) (Cohen et al., 2020; Pitts, Martínez, & Hillyard, 2012; Pitts, Padwal, Fennelly, Martínez, & Hillyard, 2014). This could also provide an explanation for the fact that we did find enhanced late-stage T1 representation, but no differences in the T1-evoked P3b between blink and no-blink trials in the same time period.

Of further note, previous ERP studies did not observe differences in T1 processing between T2 seen vs. blink trials until after 300ms. Yet, we found that the neural representation of T1 was enhanced also already at the early processing stage (150-250ms). Univariate ERP analyses are less sensitive towards changes in the pattern of activity across the scalp, which could explain this discrepancy in findings. However, it must be noted that the boost and bounce model assumes that it only takes about 100ms for the bulk of the attentional feedback to modulate the sensory representation of the evoking stimulus (Olivers & Meeter, 2008). Yet, our early T1 effect occurred after 100ms. A recent intracranial EEG study did observe a very early difference in T1 processing (Slagter et al., 2017). That is, only in T2 blink trials did T1 induce a very early (~80ms) increase in alpha/low beta activity in the ventral striatum, also suggestive of differences in early T1 processing, albeit at the subcortical level, which conceivably cannot be picked up with scalp EEG (Cohen, Cavanagh, & Slagter, 2011). Animal studies have shown similar short-latency striatal responses to salient stimuli and suggest that they may reflect a signal to frontal regions to orient attention to enhance the visual representation of a potentially relevant stimulus (Overton, Vautrelle, & Redgrave, 2014). This fits with proposals that the basal ganglia play a critical role in gating information into working memory (Hazy, Frank, & O’Reilly, 2006) and could explain the relatively “late” modulation of T1 processing observed at the scalp level in our study.

The attentional blink was also associated with differences in late-stage D1 representation. This finding could suggest that when D1 is treated like a target (i.e., is ‘spared’), as indicated by enhanced late stage decoding, T2 is also spared (i.e., seen). If true, this would critically suggest that at least some portion of T2 seen trials reflects the well-known phenomenon of extended sparing (Di Lollo et al., 2015). However, in the absence of any D1 report data, this possibility remains speculative. Future studies are necessary to replicate and determine the functional significance of our D1 effect and to replicate the here observed relationship between early T1 representational strength and the attentional blink.

It is worth noting that our pattern classifiers were likely not optimal for uncovering a wider range of processes linked to conscious access, as they were specifically sensitive to identity-specific features of target numbers and distractor letters, and as our decoding results suggested, were less revealing of more generic late encoding and working memory processes (>600ms). The AB has also been associated with relatively early differences in T2 processing, within ~200 to 300ms, as for example captured in the N2 (Sergent et al., 2005) and the N2pc (Akyürek, Leszczyński, & Schubö, 2010). Our classifiers may not have picked up on AB-related differences in T2 processing that are generic (i.e., unaffected by number identity). Arguably, our MVPA classifiers were also less sensitive to potential generic modulations of neural response gain. Selection-related boost/bounce feedback is presumably location-specific, boosting processing of stimuli presented at the same spatial location as the feedback-eliciting stimulus (Olivers & Meeter, 2008). This would suggest that the mechanism by which selection-related feedback affects subsequent processing could be similar to that of spatial attention, which has been shown to modulate neural population responses by affecting their response gain, as opposed to sharpening neuronal tuning to stimulus features (e.g. David, Hayden, Mazer, & Gallant, 2008; Fang, Becker, & Liu, 2019; Ling, Liu, & Carrasco, 2009; Williford & Maunsell, 2006). Indeed, a recent study using EEG and forward encoding modelling found that T2 selection was associated with an increase in gain, not the precision of its neural representation, suggesting that temporal attention works in a similar manner as spatial attention (Tang et al., 2020). Yet, this study unexpectedly did not find any differences in T1 or D1 representations between T2 seen and unseen trials. One notable difference between the current study and the study by Tang et al. (2020) is that the latter study examined changes in the representation of a stimulus feature (orientation) that did not dissociate targets from distractors, as target and distractors were defined by spatial frequency. As we decoded features that identified targets and distractors, it is possible that we were more sensitive to picking up effects of feature attention on sensory representations of T1 and D1. Given these opposing results, future studies that can also measure changes in the sharpness of location representations, are necessary to determine how spatial and feature attention may jointly or independently affect conscious access.

An unexpected aspect of our findings was the absence of differences in the strength of early-stage and late-stage T2 identity-specific neural representations between trials in which T2 was seen versus not seen. Previous fMRI studies have shown enhanced T2 processing in T2-seen trials, in frontal and parietal areas as well as in low-level visual areas, such as the primary visual cortex (Hein, Alink, Kleinschmidt, & Müller, 2009; Slagter, Johnstone, Beets, & Davidson, 2010; Williams, Visser, Cunnington, & Mattingley, 2008). Using EEG, Tang et al. (2020), as noted above, also observed differences in early T2 orientation representation between blink and no-blink trials, within 100-150ms post-T2. Yet, here, while we could decode T2 number identity peaking around 170 ms, we could do so equally well in T2 blink and no-blink trials. Also, late-stage T2 decoding was close to chance level regardless of T2 report. One possibility is that in a context with multiple targets, representational codes of later targets become more variable in latency or in the format (e.g., visual, phonological) in which a target is encoded or maintained, which would thus render robust classification of T2 difficult. For example, participants might have later relied on phonological representations to perform serial order recall of multiple targets in the main attention task, while the simpler localizer task could have been solved by relying on perceptual or semantic representations (Nishiyama, 2020). Several EEG studies have also shown that the latency of T2 processing is more variable during the time window of the AB (Chennu, Craston, Wyble, & Bowman, 2009; Slagter, Lutz, Greischar, Nieuwenhuis, & Davidson, 2009). Thus, variability in the latency or in the format of T2 representation may have hampered our decoding efforts. It is also possible that the inability to decode T2 at later stages is (in part) due to a rapid transformation of its representation into an activity-silent neural state. In the context of working memory, it has been shown that transiently unattended items in working memory (because another item in working memory is prioritized or attended) are no longer represented in the pattern of neural activity, but are hidden, in that they can be retrieved using an impulse stimulus or ‘ping’ (Wolff, Jochim, Akyürek, & Stokes, 2017). Activity-silent representations could more generally provide an explanation for our relative poor late-stage decoding in the attention task (in which multiple targets had to be maintained in working memory) compared to the localizer task (in which only one target had to be maintained in working memory). Lastly, the at-chance late-stage T2 identity decoding may also reflect the selective sensitivity of our classifiers to identity-specific information. In fact, using univariate analyses, we replicated the common finding of a larger T2-evoked P3b by seen compared to unseen T2s, indicative of access-related differences in late-stage T2 processing. It is of note in this regard that classifiers trained to decode whether a T2 was seen or unseen, irrespective of its identity, revealed clusters of significant decoding scores for over 900 ms after T2 presentation, confirming that the neural signal contained information related to T2 conscious access throughout the trial. This analysis also identified differences in neural activity patterns well before T2 presentation. While some of these differences likely reflect attentional blink-related differences in T1 and/or D1 processing, the pattern of scalp EEG activity already predicted conscious T2 access well before T1 was presented. This finding suggests that baseline fluctuations in neural excitability and attentional state or in temporal expectations across trials can shape the likelihood of conscious access to a significant extent, in line with previous work (Iemi et al., 2017; Kranczioch et al., 2007; Mathewson, Gratton, Fabiani, Beck, & Ro, 2009, Pincham & Szucs, 2012). Our data thus also indicate that the attentional blink is likely determined by multiple factors (e.g. Lindh, Sligte, Assecondi, Shapiro, & Charest, 2019).

To conclude, our findings do not support the notion that top-down modulation of early-stage visual representations is the major determinant of conscious access in rapidly changing input conditions as in the RSVP attention task. We did not find evidence for a rapid attentional gating mechanism that modulated early representational dynamics preceding conscious access, as proposed by the boost and bounce theory. The attentional blink was associated with differences in T1 and late D1 neural representation, and in pre-T1 activity patterns, highlighting the complex and multifaceted nature of processes determining conscious access and informing theories of attention and consciousness.

## Acknowledgements

This work was supported by a grant from the Psychology Research Institute of the University of Amsterdam.

## References

Akyürek, E. G., Leszczyński, M., & Schubö, A. (2010). The temporal locus of the interaction between working memory consolidation and the attentional blink. Psychophysiology, 47(6), 1134–1141. https://doi.org/10.1111/j.1469-8986.2010.01033.x

Bishop, M. C. (2006). Pattern Recognition and Machine Learning. The Ecstatic and the Archaic: An Analytical Psychological Inquiry. Springer. https://doi.org/10.4324/9780203733332

Bowman, H., Wyble, B., Chennu, S., & Craston, P. (2008). A reciprocal relationship between bottom-up trace strength and the attentional blink bottleneck: Relating the LC-NE and ST2 models. Brain Research, 1202, 25–42. https://doi.org/10.1016/j.brainres.2007.06.035

Bowman, Howard, & Wyble, B. (2007). The Simultaneous Type, Serial Token Model of Temporal Attention and Working Memory, 114(1), 38–70. https://doi.org/10.1037/0033-295X.114.1.38

Carlson, T., Tovar, D. A., Alink, A., & Kriegeskorte, N. (2013). Representational dynamics of object vision: The first 1000 ms. Journal of Vision, 13(10), 1–19. https://doi.org/10.1167/13.10.1

Carreiras, M., Quiñones, I., Hernández-Cabrera, J. A., & Duñabeitia, J. A. (2015). Orthographic coding: Brain activation for letters, symbols, and digits. Cerebral Cortex, 25(12), 4748–4760. https://doi.org/10.1093/cercor/bhu163

Chennu, S., Craston, P., Wyble, B., & Bowman, H. (2009). Attention increases the temporal precision of conscious perception: Verifying the neural-ST2 model. PLoS Computational Biology, 5(11). https://doi.org/10.1371/journal.pcbi.1000576

Chun, M. M., & Potter, M. C. (1995). A Two-Stage Model for Multiple Target Detection in Rapid Serial Visual Presentation. Journal of Experimental Psychology: Human Perception and Performance, 21(1), 109–127. https://doi.org/10.1037/0096-1523.21.1.109

Cohen, M. A., Ortego, K., Kyroudis, A., & Pitts, M. (2020). Distinguishing the neural correlates of perceptual awareness and post-perceptual processing. The Journal of Neuroscience, (May), JN-RM-0120-20. https://doi.org/10.1523/jneurosci.0120-20.2020

Cohen, M. X., Cavanagh, J. F., & Slagter, H. A. (2011). Event-related potential activity in the basal ganglia differentiates rewards from nonrewards: Temporospatial principal components analysis and source localization of the feedback negativity: Commentary. Human Brain Mapping, 32(12), 2270–2271. https://doi.org/10.1002/hbm.21358

Craston, P., Wyble, B., Chennu, S., & Bowman, H. (2009). The attentional blink reveals serial working memory encoding: Evidence from virtual and human event-related potentials. Journal of Cognitive Neuroscience, 21(3), 550–566. https://doi.org/10.1162/jocn.2009.21036

David, S. V., Hayden, B. Y., Mazer, J. A., & Gallant, J. L. (2008). Attention to Stimulus Features Shifts Spectral Tuning of V4 Neurons during Natural Vision. Neuron, 59(3), 509–521. https://doi.org/10.1016/j.neuron.2008.07.001

Dehaene, S., & Changeux, J. P. (2011). Experimental and Theoretical Approaches to Conscious Processing. Neuron, 70(2), 200–227. https://doi.org/10.1016/j.neuron.2011.03.018

Dehaene, S., Changeux, J. P., Naccache, L., Sackur, J., & Sergent, C. (2006). Conscious, preconscious, and subliminal processing: a testable taxonomy. Trends in Cognitive Sciences, 10(5), 204–211. https://doi.org/10.1016/j.tics.2006.03.007

Dehaene, S., Charles, L., King, J. R., & Marti, S. (2014). Toward a computational theory of conscious processing. Current Opinion in Neurobiology, 25(1947), 76–84. https://doi.org/10.1016/j.conb.2013.12.005

Dell’Acqua, R., Doro, M., Dux, P. E., & Losier, T. (2016). Enhanced frontal activation underlies sparing from the attentional blink: Evidence from human electrophysiology, 53, 623–633. https://doi.org/10.1111/psyp.12618

Derda, M., Koculak, M., Windey, B., Gociewicz, K., Wierzchoń, M., Cleeremans, A., & Binder, M. (2019). The role of levels of processing in disentangling the ERP signatures of conscious visual processing. Consciousness and Cognition, 73(June), 1–12. https://doi.org/10.1016/j.concog.2019.102767

Di Lollo, V., Kawahara, J., Ghorashi, S. M. S., & Enns, J. T. (2005). The attentional blink : Resource depletion or temporary loss of control? Psychological Research, 69, 191–200. https://doi.org/10.1007/s00426-004-0173-x

Dux, P. E., & Marois, R. (2009). The attentional blink: A review of data and theory. Attention, Perception, and Psychophysics, 71(8), 1683–1700. https://doi.org/10.3758/APP.71.8.1683

Fahrenfort, J. J., Leeuwen, J. Van, Olivers, C. N. L., & Hogendoorn, H. (2017). Perceptual integration without conscious access. Proceedings of the National Academy of Sciences, (201617268). https://doi.org/10.1073/pnas.1617268114

Fang, M. W. H., Becker, M. W., & Liu, T. (2019). Attention to colors induces surround suppression at category boundaries. Scientific Reports, 9(1), 1–13. https://doi.org/10.1038/s41598-018-37610-7

Fawcett, T. (2006). An introduction to ROC analysis. Pattern Recognition Letters, 27(8), 861–874. https://doi.org/10.1016/j.patrec.2005.10.010

Gramfort, A., Luessi, M., Larson, E., Engemann, D. A., Strohmeier, D., Brodbeck, C., … Hämäläinen, M. S. (2014). MNE software for processing MEG and EEG data. NeuroImage, 86, 446–460. https://doi.org/10.1016/j.neuroimage.2013.10.027

Grootswagers, T., Robinson, A. K., & Carlson, T. A. (2019). The representational dynamics of visual objects in rapid serial visual processing streams. NeuroImage, 188(January), 668–679. https://doi.org/10.1016/j.neuroimage.2018.12.046

Grootswagers, T., Wardle, G. S., & Carlson, A. T. (2017). Decoding Dynamic Brain Patterns from Evoked Responses: A Tutorial on Multivariate Pattern Analysis Applied to Time Series Neuroimaging Data. Journal of Cognitive Neuroscience, 29(4), 677–697. https://doi.org/10.1162/jocn

Hazy, T. E., Frank, M. J., & O’Reilly, R. C. (2006). Banishing the homunculus: Making working memory work. Neuroscience, 139(1), 105–118. https://doi.org/10.1016/j.neuroscience.2005.04.067

Hein, G., Alink, A., Kleinschmidt, A., & Müller, N. G. (2009). The attentional blink modulates activity in the early visual cortex. Journal of Cognitive Neuroscience, 21(1), 197–206. https://doi.org/10.1162/jocn.2008.21026

Iemi, L., Chaumon, M., Crouzet, S. M., & Busch, N. A. (2017). Spontaneous neural oscillations bias perception by modulating baseline excitability. Journal of Neuroscience, 37(4), 807–819. https://doi.org/10.1523/JNEUROSCI.1432-16.2016

Jolicœur, P., & Dell’Acqua, R. (1998). The Demonstration of Short-Term Consolidation, 202(36), 138–202.

Kaiser, D., Oosterhof, N. N., & Peelen, M. V. (2016). The neural dynamics of attentional selection in natural scenes. Journal of Neuroscience, 36(41), 10522–10528. https://doi.org/10.1523/JNEUROSCI.1385-16.2016

Kawahara, J. I., Kumada, T., & Di Lollo, V. (2006). The attentional blink is governed by a temporary loss of control. Psychonomic Bulletin and Review, 13(5), 886–890. https://doi.org/10.3758/BF03194014

Kihara, K., & Kawahara, J. (2008). Electrophysiological evidence for independent consolidation of multiple targets, 19(15), 3–6. https://doi.org/10.1097/WNR.0b013e32830fe4e8

King, J.-R., Pescetelli, N., & Dehaene, S. (2016). Brain Mechanisms Underlying the Brief Maintenance of Seen and Unseen Sensory Information Article Brain Mechanisms Underlying the Brief Maintenance of Seen and Unseen Sensory Information, 1122–1134. https://doi.org/10.1016/j.neuron.2016.10.051

King, J. R., & Dehaene, S. (2014). Characterizing the dynamics of mental representations: The temporal generalization method. Trends in Cognitive Sciences, 18(4), 203–210. https://doi.org/10.1016/j.tics.2014.01.002

Kranczioch, C., Debener, S., Maye, A., & Engel, A. K. (2007). Temporal dynamics of access to consciousness in the attentional blink. NeuroImage, 37(3), 947–955. https://doi.org/10.1016/j.neuroimage.2007.05.044

Lagroix, H. E. P., Spalek, T. M., Wyble, B., Jannati, A., & Di Lollo, V. (2012). The root cause of the attentional blink: First-target processing or disruption of input control? Attention, Perception, and Psychophysics, 74(8), 1606–1622. https://doi.org/10.3758/s13414-012-0361-5

Lamme, V. a F., & Roelfsema, P. R. (2000). The distinct modes of vision offered by feedforward and recurrent processing. Trends in Neurosciences, 23(11), 571–579. https://doi.org/10.1016/S0166-2236(00)01657-X

Lindh, D., Sligte, I. G., Assecondi, S., Shapiro, K. L., & Charest, I. (2019). Conscious perception of natural images is constrained by category-related visual features. Nature Communications, 10(1), 1–9. https://doi.org/10.1038/s41467-019-12135-3

Ling, S., Liu, T., & Carrasco, M. (2009). How spatial and feature-based attention affect the gain and tuning of population responses. Vision Research, 49(10), 1194–1204. https://doi.org/10.1016/j.visres.2008.05.025

Lunau, R., & Olivers, C. N. L. (2010). The attentional blink and lag 1 sparing are nonspatial. Attention, Perception, & Psychophysics, 72(2), 317–325. https://doi.org/10.3758/APP

Maris, E., & Oostenveld, R. (2007). Nonparametric statistical testing of EEG- and MEG-data, 164(1), 177–190. https://doi.org/10.1016/j.jneumeth.2007.03.024

Martens, S., Elmallah, K., London, R., & Johnson, A. (2006). Cuing and stimulus probability effects on the P3 and the AB. Acta Psychologica, 123(3), 204–218. https://doi.org/10.1016/j.actpsy.2006.01.001

Martens, S., Munneke, J., Smid, H., & Johnson, A. (2006). Quick minds don’t blink: Electrophysiological correlates of individual differences in attentional selection. Journal of Cognitive Neuroscience, 18(9), 1423–1438. https://doi.org/10.1162/jocn.2006.18.9.1423

Marti, S., & Dehaene, S. (2017). Discrete and continuous mechanisms of temporal selection in rapid visual streams. Nature Communications, 8(1). https://doi.org/10.1038/s41467-017-02079-x

Marti, S., Sigman, M., & Dehaene, S. (2012). A shared cortical bottleneck underlying attentional blink and psychological refractory period. NeuroImage, 59(3), 2883–2898. https://doi.org/10.1016/j.neuroimage.2011.09.063

Martin, J. G., Cox, P. H., Scholl, C. A., & Riesenhuber, M. (2019). A crash in visual processing: Interference between feedforward and feedback of successive targets limits detection and categorization, 19, 1–21.

Mathewson, K. E., Gratton, G., Fabiani, M., Beck, D. M., & Ro, T. (2009). To See or Not to See: Prestimulus α Phase Predicts Visual Awareness. The Journal of Neuroscience, 29(9), 2725–2732. https://doi.org/10.1523/JNEUROSCI.3963-08.2009

Meijs, E. L., Mostert, P., Slagter, H. A., de Lange, F. P., & van Gaal, S. (2019). Exploring the role of expectations and stimulus relevance on stimulus-specific neural representations and conscious report. Neuroscience of Consciousness, 2019(1), 1–12. https://doi.org/10.1093/nc/niz011

Myerson, J., Green, L., & Warusawitharana, M. (2001). Area Under the Curve As a Measure of Discounting. Journal of the Experimental Analysis of Behavior, 76(2), 235–243. https://doi.org/10.1901/jeab.2001.76-235

Niedeggen, M., Hesselmann, G., Sahraie, A., Milders, M., & Blakemore, C. (2004). Probing the prerequisites for motion blindness. Journal of Cognitive Neuroscience, 16(4), 584–597. https://doi.org/10.1162/089892904323057317

Nishiyama, R. (2020). Adaptive use of semantic representations and phonological representations in verbal memory maintenance. Journal of Memory and Language, 111(December 2019), 104084. https://doi.org/10.1016/j.jml.2019.104084

Olivers, C. N. L. (2012). The attentional bost and the attentional blink. https://doi.org/10.1093/acprof

Olivers, C. N. L., Hilkenmeier, F., & Scharlau, I. (2011). Prior entry explains order reversals in the attentional blink. Attention, Perception, and Psychophysics, 73(1), 53–67. https://doi.org/10.3758/s13414-010-0004-7

Olivers, C. N. L., & Meeter, M. (2008). A Boost and Bounce Theory of Temporal Attention. Psychological Review, 115(4), 836–863. https://doi.org/10.1037/a0013395

Olivers, C. N. L., Van Der Stigchel, S., & Hulleman, J. (2007). Spreading the sparing: Against a limited-capacity account of the attentional blink. Psychological Research, 71(2), 126–139. https://doi.org/10.1007/s00426-005-0029-z

Overton, P. G., Vautrelle, N., & Redgrave, P. (2014). Sensory regulation of dopaminergic cell activity: Phenomenology, circuitry and function. Neuroscience, 282, 1–12. https://doi.org/10.1016/j.neuroscience.2014.01.023

Pedregosa, F., Weiss, R., & Brucher, M. (2011). Scikit-learn: Machine Learning in Python, 12, 2825–2830.

Pincham, H. L., & Szucs, D. (2012). Conscious access is linked to ongoing brain state: Electrophysiological evidence from the attentional blink. Cerebral Cortex, 22(10), 2346–2353. https://doi.org/10.1093/cercor/bhr314

Pitts, M. A., Martínez, A., & Hillyard, S. A. (2012). Visual processing of contour patterns under conditions of inattentional blindness. Journal of Cognitive Neuroscience, 24(2), 287–303. https://doi.org/10.1162/jocn_a_00111

Pitts, M. A., Padwal, J., Fennelly, D., Martínez, A., & Hillyard, S. A. (2014). Gamma band activity and the P3 reflect post-perceptual processes, not visual awareness. NeuroImage, 101, 337–350. https://doi.org/10.1016/j.neuroimage.2014.07.024

Raymond, J. E., Shapiro, K. L., & Arnell, K. M. (1992). Temporary Suppression of Visual Processing in an RSVP Task: An Attentional Blink? Journal of Experimental Psychology: Human Perception and Performance. https://doi.org/10.1037/0096-1523.18.3.849

Sergent, C., Baillet, S., & Dehaene, S. (2005). Timing of the brain events underlying access to consciousness during the attentional blink. Nature Neuroscience, 8(10), 1391–1400. https://doi.org/10.1038/nn1549

Shanks, D. R. (2017). Regressive research: The pitfalls of post hoc data selection in the study of unconscious mental processes. Psychonomic Bulletin and Review, 24(3), 752–775. https://doi.org/10.3758/s13423-016-1170-y

Shapiro, K., Schmitz, F., Martens, S., Hommel, B., & Schnitzler, A. (2006). Resource sharing in the attentional blink. NeuroReport, 17(2), 163–166. https://doi.org/10.1097/01.wnr.0000195670.37892.1a

Shimozaki, S. S., Chen, K. Y., Abbey, C. K., & Eckstein, M. P. (2007). The temporal dynamics of selective attention of the visual periphery as measured by classification images. Journal of Vision, 7(12), 1–20. https://doi.org/10.1167/7.12.10

Sigman, M., & Dehaene, S. (2008). Brain mechanisms of serial and parallel processing during dual-task performance. Journal of Neuroscience, 28(30), 7585–7598. https://doi.org/10.1523/JNEUROSCI.0948-08.2008

Slagter, H. A., Johnstone, T., Beets, I. A. M., & Davidson, R. J. (2010). Neural competition for conscious representation across time: An fMRI study. PLoS ONE, 5(5), 1–10. https://doi.org/10.1371/journal.pone.0010556

Slagter, H. A., Lutz, A., Greischar, L. L., Francis, A. D., Nieuwenhuis, S., Davis, J. M., & Davidson, R. J. (2007). Mental training affects distribution of limited brain resources. PLoS Biology, 5(6), 1228–1235. https://doi.org/10.1371/journal.pbio.0050138

Slagter, H. A., Lutz, A., Greischar, L. L., Nieuwenhuis, S., & Davidson, R. J. (2009). Theta phase synchrony and conscious target perception: Impact of intensive mental training. Journal of Cognitive Neuroscience, 21(8), 1536–1549. https://doi.org/10.1162/jocn.2009.21125

Slagter, H. A., Mazaheri, A., Reteig, L. C., Smolders, R., Figee, M., Mantione, M., … Denys, D. (2017). Contributions of the ventral striatum to conscious perception: An intracranial EEG study of the attentional blink. Journal of Neuroscience, 37(5), 1081–1089. https://doi.org/10.1523/JNEUROSCI.2282-16.2016

Tang, M. F., Ford, L., Arabzadeh, E., Enns, J. T., Visser, T. A. W., & Mattingley, J. B. (2020). Neural dynamics of the attentional blink revealed by encoding orientation selectivity during rapid visual presentation. Nature Communications, 11(1), 1–14. https://doi.org/10.1038/s41467-019-14107-z

van Moorselaar, D., & Slagter, H. A. (2019). Learning What Is Irrelevant or Relevant: Expectations Facilitate Distractor Inhibition and Target Facilitation through Distinct Neural Mechanisms. The Journal of Neuroscience : The Official Journal of the Society for Neuroscience, 39(35), 6953–6967. https://doi.org/10.1523/JNEUROSCI.0593-19.2019

Vogel, E. K., Luck, S. J., & Shapiro, K. L. (1998). Electrophysiological Evidence for a Postperceptual Locus of Suppression during the Attentional Blink. Journal of Experimental Psychology: Human Perception and Performance, 24(6), 1656–1674. https://doi.org/10.1037/0096-1523.24.6.1656

Wagenmakers, E., Love, J., Maarten, L., Tahira, M., Ly, A., Verhagen, J., … Morey, R. D. (2018). Bayesian inference for psychology. Part II: Example applications with JASP. Psychonomic Bulletin & Review, 25(1), 58–76. https://doi.org/10.3758/s13423-017-1323-7

Weaver, M. D., Fahrenfort, J. J., Belopolsky, A., & Van Gaal, S. (2019). Independent neural activity patterns for sensory-and confidence-based information maintenance during category-selective visual processing. ENeuro, 6(1). https://doi.org/10.1523/ENEURO.0268-18.2018

Williams, M. A., Visser, T. A. W., Cunnington, R., & Mattingley, J. B. (2008). Attenuation of neural responses in primary visual cortex during the attentional blink. Journal of Neuroscience, 28(39), 9890–9894. https://doi.org/10.1523/JNEUROSCI.3057-08.2008

Williford, T., & Maunsell, J. H. R. (2006). Effects of spatial attention on contrast response functions in macaque area V4. Journal of Neurophysiology, 96(1), 40–54. https://doi.org/10.1152/jn.01207.2005

Wyble, B., Bowman, H., & Nieuwenstein, M. (2009). The Attentional Blink Provides Episodic Distinctiveness: Sparing at a Cost. Journal of Experimental Psychology: Human Perception and Performance, 35(3), 787–807. https://doi.org/10.1037/a0013902

Wyble, B., Bowman, H., & Potter, M. C. (2009). Categorically Defined Targets Trigger Spatiotemporal Visual Attention. Journal of Experimental Psychology: Human Perception and Performance, 35(2), 324–337. https://doi.org/10.1037/a0013903

